# A cis-regulatory change underlying the motor neuron-specific loss of terminal selector gene expression in immotile tunicate larvae

**DOI:** 10.1101/567719

**Authors:** Elijah K. Lowe, Claudia Racioppi, Nadine Peyriéras, Filomena Ristoratore, Lionel Christiaen, Billie J. Swalla, Alberto Stolfi

## Abstract

The evolutionary history of animal body plans cannot be fully reconstructed without considering the roles of both novelties and losses. Some of the more remarkable examples of massively parallel evolutionary losses in animals comes from many species in the tunicate genus *Molgula* that have independently lost the swimming larva and instead develop as tail-less, immotile larvae that bypass the period of swimming and dispersal observed in other tunicates, marine invertebrate chordates that alternate between motile larval and sessile adult life cycle stages. The larvae of *Molgula occulta* and other tail-less species do not fully develop structures that are essential for swimming behavior, including notochord, tail muscles, and otolith, and loss-of-function mutations have been identified in various genes required for the differentiation of these tissues. However, little is known about the extent of development of the larval nervous system in *M. occulta*. While differentiated neurons might in principle be entirely dispensable to the non-swimming larva, the adult has a fully functional nervous system like any other tunicate. To further investigate this conundrum, we studied the specification and patterning of the *M. occulta* Motor Ganglion, which is the key central nervous system compartment that drives the motor movements of swimming tunicate larvae. We found that the expression patterns of important regulators of MG neuron subtype specification are highly conserved during the development of the non-swimming larvae of *M. occulta*, suggesting that the gene networks regulating their expression are largely intact in this species, despite the loss of swimming ability. However, we identified a *M. occulta*-specific reduction in expression of the important motor neuron terminal selector gene *Ebf (Collier/Olf/EBF or COE)* in the Motor Ganglion. Although *M. occulta Ebf* is predicted to encode a fully functional protein, its expression was reduced in developing motor neurons when compared to species with swimming larvae, which was corroborated by measuring allele-specific expression of *Ebf* in interspecific hybrid embryos produced by crossing *M. occulta* with the closely related swimming species *M. oculata*. Comparative reporter construct experiments also revealed a specific *cis*-regulatory sequence change that underlies the reduced expression of *M. occulta Ebf* in motor neurons, but not in other tissues and cell types. This points to a potential mechanism for arresting larval motor neuron differentiation in the non-swimming larvae of this species.

## Introduction

The evolution of animal body plans has occurred not only through evolutionary novelties, but also through losses both subtle and catastrophic (Albalat and Cañestro, 2016). Although they are defined primarily by the absence of structures, cell types, or genes, the evolutionary loss of these various units of selection can help illuminate their functions. For instance, identifying traits that are retained in one species but lost in a closely related species can reveal which are likely to be under purifying selection and provide a measure of their relative adaptive value under certain conditions, e.g. loss of vision in cave-dwelling organisms (Porter et al., 2003).

Some of the most prominent examples of extensive evolutionary losses come from the tunicates (Denoeud et al., 2010; Huber et al., 2000), marine filter-feeding organisms characterized by a protective layer, or “tunic”, made mostly of cellulose (Sasakura, 2018). Phylogenomic analyses suggest that vertebrates are the sister group to the tunicates (Bourlat et al., 2006), and most tunicates have a distinct dispersal phase carried out by swimming “tadpole” type larvae that bear the typical chordate body plan: a rigid notochord flanked by striated paraxial muscles, controlled by a dorsal central nervous system. Some tunicate groups, such as the pelagic salps and pyrosomes, have completely lost this larval phase, while in other groups the evolutionary loss of the swimming larva is an ongoing process, affecting specific aspects of larval development and behavior. This phenomenon is particularly prominent in molgulid tunicates, in which many species have independently lost larval structures important for swimming (Hadfield et al., 1995; Huber et al., 2000). The best studied of these is *Molgula occulta*, which gives rise to anural (tail-less), non-swimming larvae that metamorphose into juveniles without going through the dispersal period of active swimming observed in urodele (tailed) species (Berrill, 1931).

Underlying this radically divergent, non-swimming larval form is the loss of morphogenesis and differentiation of certain cell types that are dispensable for non-swimming larva, like the notochord, tail muscles, or pigmented sensory organs (Jeffery and Swalla, 1991; Swalla and Jeffery, 1990, 1992). For instance, *M. occulta* larva muscle cells do not differentiate and the species has even lost certain genes encoding proteins specifically required for muscle function, like muscle actin (Kusakabe et al., 1996). Furthermore, *M. occulta* lack a pigmented otolith cell, important for gravity sensing in swimming larvae (Jiang et al., 2005; Tsuda et al., 2003), due to the loss of genes encoding functional enzymes required for melanogenesis like Tyrosinase and Tyrosinase-related protein (Tyrp)(Racioppi et al., 2017). Over 17 species with non-swimming larvae have been described in *Molgula*, and all 31 species in the closely related molgulid genera *Eugyra* and *Bostrichobranchus* appear to have non-swimming larvae. The vast majority of molgulids (>150 species) have not been studied at the larval stage, leaving the possibility that many more non-swimming larvae have independently evolved within the clade (Maliska et al., 2013; Shenkar et al., 2019).

Little is known about the extent of nervous system development in the larvae of *M. occulta* and other species with non-swimming larvae. The typical tunicate larva has a minimal nervous system dedicated to controlling its swimming and settlement behavior in response to sensory cues such as light, gravity, and mechanical stimuli (Jiang et al., 2005; Rudolf et al., 2018; Salas et al., 2018; Zega et al., 2006). The larval nervous system of the tunicate *Ciona intestinalis* has been completely mapped, revealing 177 neurons in a dorsal central nervous system (CNS) and 54 peripheral sensory neurons distributed throughout the epidermis (Ryan et al., 2016, 2018; Ryan and Meinertzhagen, 2019). With 231 total neurons, the *Ciona* larval nervous system is one of the smallest known in all of metazoa, and thus an intriguing model for the study of chordate-specific principles of neuronal function and development (Nishino, 2018).

Given the evolutionary loss of other structures important for swimming (notochord, muscles, otolith), we asked whether any neurodevelopmental processes have been lost during the evolution of *M. occulta*. To this end, we surveyed the development of the nervous system in *M. occulta* embryos, using species with swimming larvae as a basis for comparison. Using *in situ* hybridization, RNAseq data, and cross-species transgenic assays, we report that neurodevelopmental gene expression and patterning is unexpectedly conserved in *M. occulta* larvae. Most notably, the Motor Ganglion (MG), the CNS structure that controls the swimming movements of tailed larvae, is also found in tailless larvae (Nishino et al., 2010). However, we uncovered specific *cis*-regulatory mutations that might underlie the reduced transcriptional activity of the key cholinergic neuron terminal selector gene *Ebf (Kratsios et al., 2012)* in larval motor neurons. However, unlike *Tyrosinase* or *Tyrp*, the *Ebf* gene has not become pseudogenized in *M. occulta*, likely due to its requirement for the specification of other neurons and cell types that are still important for the complete life cycle of this species.

## Methods

### *M. occulta* and *M. oculata* embryo collection

Gonads were dissected from gravid *M. occulta* and *M. oculata* adults were collected in August (the only time of the year when gravid individuals can be found there) at Station Biologique in Roscoff, France. *Molgula occidentalis* adults were collected and shipped by Gulf Specimen Marine Lab (Panacea, FL). Eggs from dissected gonads were fertilized *in vitro*, dechorionated, and fixed as previously described (Stolfi et al., 2014b).

### mRNA probe synthesis, *in situ* hybridization, and immunostaining

Templates for mRNA *in situ* hybridization probes were cloned by PCR or SMARTer 3’/5’-RACE (Clontech) from cDNA or genomic DNA (see **Supplemental File 1** for details). *In vitro* transcription of labeled probes was and two-color fluorescent *in situ* hybridization were performed as previously described (Ikuta and Saiga, 2007; Stolfi et al., 2014b). To detect cilia, larvae were incubated with anti-acetylated α-tubulin primary antibody (clone 6-11B-1, Sigma) and AlexaFluor-488 secondary antibody (ThermoFisher) as previously described (Pennati et al., 2015). Cell outlines were counterstained with phalloidin AlexFluor-546 conjugate (ThermoFisher) incubated 1:50 for at least 2 hours prior to final washes, and nuclei were labeled with DAPI during the final wash. Embryos were imaged on upright and inverted epifluorescence microscopes (Leica), or TCS SP5 AOBS and TCS SP8 X scanning point confocal systems (Leica).

### Hybrid embryo allele-specific differential expression analyses

*M. occulta* and *M. oculata* genomes (Stolfi et al., 2014b) were improved using Redundans (Pryszcz and Gabaldón, 2016), reads from the initial assemblies (<u>PRJNA253689</u>) were used to improve scaffolding, which had been done to improve the *M. occidentalis* genome and gene models we previously published (Lowe and Stolfi, 2018). Additionally, the *M. oculata* genome was used as a reference to increase the scaffolding of the *M. occulta* genome. To build gene models, previously sequenced RNAseq reads (Lowe et al., 2014) were mapped using hisat2 (v2.1.0)(Kim et al., 2015), converted to BAM and sorted using SAMTools (v1.5) (Li et al., 2009) and merged with picard tools (v2.0.1) (http://broadinstitute.github.io/picard/). Merged bam files were then assembled using Trinity genome-guide command (v2.6.6)(Grabherr et al., 2011). Assembled transcripts were then translated and filtered using TransDecoder (https://github.com/TransDecoder/TransDecoder/wiki) removing any open-reading frames less than 100 amino acids. To improve on contiguity, transcripts were BLASTed against *Ciona robusta* predicted proteins from ANISEED (Brozovic et al., 2018), then scaffolded using TransPS (Liu et al., 2014). Default parameters were used for all steps and scripts can be found at https://github.com/elijahlowe/tailed. Genomes and transcriptomes used in this study can be found at https://osf.io/mj3r7/. For differential expression analysis, previously sequenced RNAseq reads from *M. occulta, M. oculata* and *M. occulta* × *M. oculata* hybrid embryos from three different stages (Fodor et al. in preparation; Lowe et al., in preparation)(Lowe et al., 2014) were mapped to the *M. occulta* or *M. oculata Ebf* gene models using Salmon v0.6.0 (Patro et al., 2015).

### Reporter construct cloning and mutagenesis

*Ebf* upstream *cis*-regulatory sequences were cloned from *M. occulta* and *M. oculata* genomic DNA (see **Supplemental File 1** for sequences). Mutations to convert *occulta* sequences to *oculata-like* sequences and vice-versa were generated by synthesizing DNA fragments (Twist Bioscience) and subcloning into the full-length reporter plasmids (see **Supplemental File 1** for sequences).

### *Ciona robusta* electroporation

*Ciona robusta* (i.e. *intestinalis* Type A) adults were collected and shipped by M-REP (San Diego, CA). Eggs were obtained from dissected adults and dechorionated, fertilized, electroporated, and fixed as previously described (Christiaen et al., 2009a, b; Stolfi et al., 2014b).

## Results and discussion

### The nervous system of *M. occulta* larvae

*M. occulta* (**Figure 1A**) and *M. oculata* (**Figure 1B**) are closely related species that occur sympatrically off the coast of Brittany, France. In spite of their close genetic similarity and their ability to form interspecific hybrids (Swalla and Jeffery, 1990), the larvae of *M. occulta* are tail-less and non-swimming (**Figure 1C**). Given its inability to swim, we asked whether the *M. occulta* larva has a functional nervous system or whether it has been partially lost in evolution like its vestigial notochord, muscles, and pigment cells. Confocal imaging of phalloidin-stained hatching *M. occulta* larvae revealed that they possess a dorsal hollow neural tube, though it is shortened along the anterior-posterior (A-P) axis due to the lack of an extended tail (**Figure 1D**). Acetylated tubulin immunohistochemistry revealed ciliated cells lining the neuropore and the shortened caudal neural tube (**Figure 1E**). In the epidermis, we found that only 4 cells at the very posterior tip of the tail bore long cilia typical of tunicate epidermal sensory cells (**Figure 1F**). Swimming larvae of tunicate species in several different families have an extended caudal nerve cored lined with numerous ciliated ependymal cells down the length of the tail, and bear numerous sensory neurons with elongated cilia, scattered throughout the epidermis of the head and tail (**Figure 1G**)(Pasini et al., 2006; Ryan et al., 2018; Torrence and Cloney, 1982; Vorontsova et al., 1997). Taken together, these data indicate that both the CNS and peripheral sensory cells of the *M. occulta* larva are only partially formed in comparison to typical solitary tunicate larvae, consistent with the hypothesis that a fully functional larval nervous system may be dispensable for this species.

**FIGURE 1 –.**
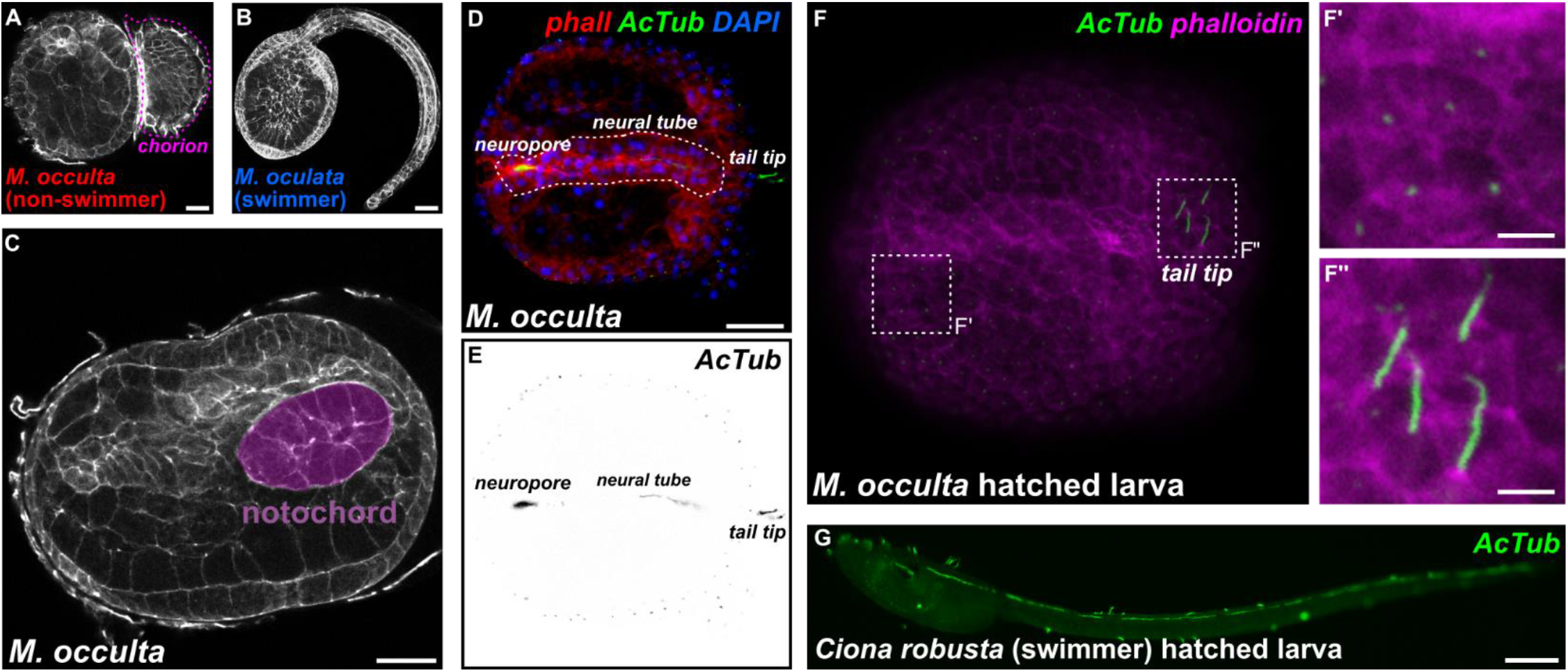
Development and morphology of the non-swimming larvae of *Molgula occulta*. **A)** Hatching larva of non-swimming species *Molgula occulta*, squeezing out of the chorion (pink dotted outline). **B)** Hatched larva of swimming species *Molgula oculata*. **C)** Hatching *M. occulta* larva showing twenty notochord cells (purple) that fail to undergo convergent extension. **D)** *M. occulta* larva stained with phalloidin conjugate (phall, red), anti-acetylated alpha tubulin antibody (AcTub, green), and DAPI (blue), showing presence of acetylated alpha tubulin-rich cilia lining the neuropore/stomodeum and neural tube lumen, and tail tip epidermal cells. **E)** Inverted monochrome image of acetylated alpha tubulin stain in D). **F)** *M. occulta* larva stained with anti-acetylated alpha tubulin antibody (AcTub, green) and phalloidin conjugate (magenta). Inset in F’ shows short cilia of epidermal cells covering the majority of the larva. Inset in F” shows longer cilia (presumably sensory) of four tail tip cells. **G)** Swimming larva of *Ciona robusta* stained with same anti-acetylated alpha tubulin antibody (AcTub, green), showing abundance of ciliated epidermal sensory cells and neural tube ependymal cells along the entire length of the larva. Scale bars in A-D: 25 μm. Scale bars in F: 5 μm. Scale bar in G: 75 μm.

To survey CNS development in *M. occulta* embryos, we performed whole-mount mRNA *in situ* hybridizations for genes that have been well characterized during CNS development in other tunicates with swimming larvae, notably *Ciona* spp., *Halocynthia roretzi*, and *Molgula occidentalis*. The first candidates we analyzed were *Celf3/4/5* (also known as *ETR-1*) *Onecut*, and *Neurogenin*, all relatively broad markers of neuronal fate in swimming tunicate larvae (**Figure 2A-C**)(D’Aniello et al., 2011; Imai et al., 2009; Lowe and Stolfi, 2018; Satou et al., 2001; Yagi and Makabe, 2001). Their mRNA *in situ* patterns clearly revealed the developing embryonic neural tube around the tailbud stage (~7.5-9.5 hours post-fertilization, hpf). Although *M. occulta* embryos are more difficult to orient given their nearly perfect spherical shape, the expression patterns formed the basic outline of the brain and MG, appearing quite conserved relative to the orthologous expression patterns in swimming species, especially *M. occidentalis* (Lowe and Stolfi, 2018).

**FIGURE 2 –.**
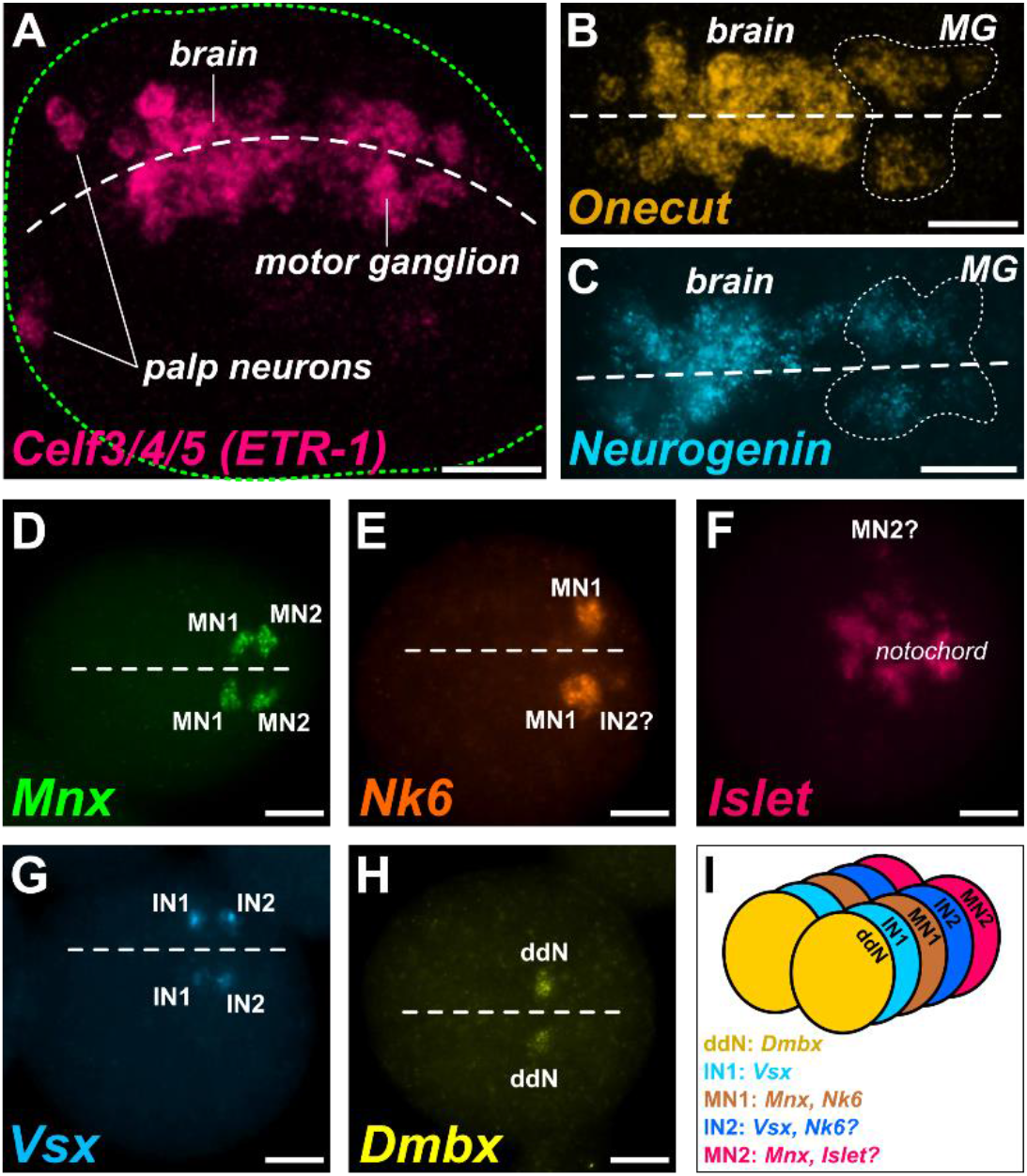
Expression patterns in the developing CNS of *Molgula occulta*. **A)** Fluorescent whole mount *in situ* hybridization (ISH) in tailbud embryos for transcripts of neural marker gene *Celf3/4/5*, also known as *ETR-1*, showing the developing nervous system including brain, motor ganglion and palps. Embryonic dorsal midline indicated by white dashed line. Embryo outline indicated by green dashed line. **B)** ISH for neural regulatory gene *Onecut*, showing brain and motor ganglion (MG)-specific expression. Onecut-expressing MG cells outlined by dotted line. **C)** ISH for proneural bHLH gene *Neurogenin*, showing brain and motor ganglion (MG)-specific expression. *Neurogenin-expressing* MG cells outlined by dotted line. **D)** ISH for motor neuron-specific marker *Mnx*, showing expression marking putative Motor Neuron 1 (MN1) and Motor Neuron 2 (MN2) bilateral pairs. **E)** ISH for *Nk6*, showing MN1-specific expression and potentially weak expression in MG Interneuron 2 (IN2) on one side. **F)** ISH for *Islet*, showing expression in disorganized notochord cells, and potentially in one MN2 cell. **G)** ISH for MG interneuron marker gene *Vsx*, showing expression in MG Interneuron 1 (IN1) and MG interneuron 2 (IN2) bilateral pairs. **H)** ISH for descending decussating neuron (ddN)-specific marker *Dmbx*. **I)** Cartoon diagram of MG neuron subtype organization in *M. occulta* based on ISH data and comparisons to the MG of swimming species, namely *Ciona robusta* and *Molgula occidentalis*. Dorsal midline in B-E and G-H indicated by dashed line. Scale bars: 25 μm

Unfortunately, we were unable to visualize *M. occulta* neurons in greater detail using fluorescent reporter plasmids, as we have done in *Ciona* and *M. occidentalis* (Lowe and Stolfi, 2018; Stolfi and Levine, 2011). This is because *M. occulta* eggs are extremely difficult to fully dechorionate and electroporate. Although we previously reported obtaining a rare electroporated *M. occulta* embryo (Stolfi et al., 2014b), our low success rate and the limited geographic range and spawning season of *M. occulta/oculata* means that we have yet to perform routine transfection of these species. Under these conditions, we proceeded with our analysis using mainly *in situ* hybridization and next-generation sequencing in *M. occulta* embryos, and heterologous reporter construct assays in *C. robusta* embryos.

### Gene expression patterns in the *M. occidentalis* Motor Ganglion

In *Ciona*, the swimming and escape-response behaviors of the larva are controlled by the MG, a central pattern generator comprised of 22 total neurons (Nishino et al., 2010; Ryan et al., 2016). Within the MG, there is a core of 8 left/right pairs of neurons that form the bulk of the synaptic connections within the MG, as well as the majority of neuromuscular synapses (Ryan et al., 2016). From now on, we will refer to the core MG neurons on only one side of the larva, with the implicit understanding that our discussion encompasses each left/right pair. We have previously characterized the core cell lineages of the MG in *Ciona* and *M. occidentalis*, documenting the highly conserved, invariant specification of each MG neuron subtype and their diagnostic marker gene expression patterns (Imai et al., 2009; Lowe and Stolfi, 2018; Stolfi and Levine, 2011; Stolfi et al., 2011). If the major function of the MG in tunicate larvae is to simply drive swimming behavior, it is likely of little adaptive value and therefore likely to be lost in a non-swimming larva like that of *M. occulta*. On the other hand, MG progenitors might also contribute to the adult nervous system, or differentiated MG neurons might be carrying out other functions beyond swimming, such as triggering metamorphosis. We therefore sought to characterize the expression patterns of candidate regulators of MG patterning in *M. occulta* embryos.

### Motor neurons MN1 and MN2

To ask whether larval motor neurons are specified in *M. occulta*, we performed *in situ* hybridization with *Mnx*, a conserved regulator of motor neuron fate (Ferrier et al., 2001) which in *Ciona* and *M. occidentalis* labels the two major motor neurons of the MG, MN1 and MN2 (Ryan et al., 2016). To our surprise, we detected exactly two left/right pairs of cells expressing *Mnx* in *M. occulta* tailbud embryos (**Figure 2D**). This staining appeared identical to *Mnx* expression in *Ciona* and *M. occidentalis*, with a gap between the two cells on either side where Interneuron 2 (IN2) should be. This suggests that *M. occulta* embryos specify both major motor neurons found in swimming species.

To possibly distinguish the specification of the two motor neuron subtypes, we did *in situ* hybridization for *Nk6* and *Islet*, markers specifically for MN1 and MN2 respectively. *Nk6* strongly labeled one or two cells on either side of the MG (**Figure 2E**), similar to its expression in *M. occidentalis* (Lowe and Stolfi, 2018). These two cells likely correspond to MN1 and IN2, based on their positions and the *M. occidentalis* pattern. In contrast, we were unable to clearly identify MN2 based on *Islet* expression, as this gene was expressed in a mass of cells in the middle of the embryo, confounding any hope of distinguishing MN2 (**Figure 2F**). This mass of cells undoubtedly corresponds to notochord cells, which do not intercalate in *M. occulta* (**Figure 1C**). In *Ciona* and *M. occidentalis, Islet* is also strongly expressed in notochord cells, but because these converge and extend in a very orderly manner, MN2 is clearly distinguishable as an Islet-expressing cell in the developing MG just dorsal to the notochord (Lowe and Stolfi, 2018; Stolfi and Levine, 2011). We were not able to visualize such clear *Islet* expression in MN2, but we conclude that MN1 and MN2 are indeed specified, based on the combined *Nk6* and *Mnx* expression patterns.

### MG Interneurons 1 and 2 (MGIN1 and MGIN2)

In the *Ciona* MG connectome, three pairs of descending interneurons are predicted to play an important role in both the rhythmicity of swimming movements and their modulation by inputs from the sensory vesicle (Kourakis et al., 2019; Ryan et al., 2016; Salas et al., 2018). Of these, the best studied are MGIN1 and MGIN2 (referred to from now on as IN1 and IN2 respectively), which flank MN1. Both are presumed excitatory interneurons that arise from the A9.30 lineage (Cole and Meinertzhagen, 2004). In *Ciona* and *M. occidentalis*, they are marked by expression of *Vsx* (Imai et al., 2009; Stolfi and Levine, 2011), the ortholog of conserved spinal cord interneuron regulator Vsx2/Chx10 (Altun-Gultekin et al., 2001; Kimura et al., 2006; Liu et al., 1994).

We found that in *M. occulta, Vsx* labels two cells on either side of the neural tube at the tailbud stage (**Figure 2G**). This is similar to the orthologous expression patterns in *Ciona* and *M. occidentalis*. In *Ciona, Vsx* is activated in IN1 and IN2 at two different time points (Stolfi and Levine, 2011). IN2 is the first interneuron to differentiate, and activates *Vsx* early, before IN1 is even born Later, after IN1 is born and specified, *Vsx* is activated in this cell. In *M. occidentalis*, we showed that *Vsx* expression appears concurrently in IN2 and in a progenitor cell that will give rise to IN1, indicating a temporal shift towards precocious activation of *Vsx* in the IN1 lineage (Lowe and Stolfi, 2018). Here, we were unable to ascertain the relative timing of *Vsx* expression and the identities of the *Vsx*-expressing cells but can conclude that IN1 and IN2 specification is conserved in *M. occulta*.

### Descending decussating neurons

The descending decussating neuron (ddN) is a highly unique neuron at the periphery of the core MG in both *Ciona* and *M. occidentalis*. It is the only descending neuron that decussates, or projects its axon contralaterally (as the name implies). Due to its position and connectivity within the *Ciona* MG, it was proposed to be the homolog of vertebrate Mauthner cells and to function in a homologous escape response pathway (Ryan et al., 2017; Takamura et al., 2010). In *Ciona* and *M. occidentalis*, the ddN is marked by expression of *Dmbx* (Lowe and Stolfi, 2018; Takahashi and Holland, 2004). In *M. occulta* tailbud embryos, *Dmbx* was found to be expressed in a single pair of cells (**Figure 2H**), likely corresponding to the ddNs or their immediate progenitors, the A11.120 cells. We conclude that the *M. occulta* larva specifies ddNs despite being physically incapable of engaging in escape response maneuvers. Taken together, these data reveal that neuronal subtypes are arrayed in the *M. occulta* MG in a manner that is identical to that previously described in swimming species such as *M. occidentalis* and *Ciona* spp. (**Figure 2I**). This suggests that, in spite of the non-swimming nature of the *M. occulta* larva, MG neuron specification and patterning have not been evolutionarily lost.

### Reduced expression of *Ebf* as a potential mechanism of impaired MG neuron differentiation

Although developmental patterning of the MG appears to be conserved in the non-swimming larva of *M. occulta*, we further pursued the hypothesis that perhaps the differentiation of specific MG neurons may not occur in *M. occulta*, similar to how the differentiation (but not the specification) of notochord, tail muscle, and otolith cells have been lost. To address this possibility, we analyzed other genes known to be involved in motor neuron differentiation. One of these is the transcription factor Ebf (also known as Collier/Olf/EBF or COE), a conserved “terminal selector” (Hobert, 2008) for cholinergic motor neurons (Kratsios et al., 2012). In *Ciona*, Ebf is required for MN2 differentiation (Stolfi et al., 2014a) and is sufficient to activate cholinergic gene expression when ectopically expressed in other cell types (Kratsios et al., 2012). When we looked at *Ebf* expression in *M. occulta*, we noticed that it appeared substantially weaker in the MG (but not in other cells) compared to its orthologous expression in *M. occidentalis*, although this assay was not quantitative (**Figure 3A,B**). Since Ebf regulates key steps between specification and differentiation in motor neurons, we hypothesized that *Ebf* expression might have been evolutionarily lost in the vestigial MG of *M. occulta*. To test this further, we turned to a more quantitative interspecies comparison of gene expression.

**FIGURE 3 –.**
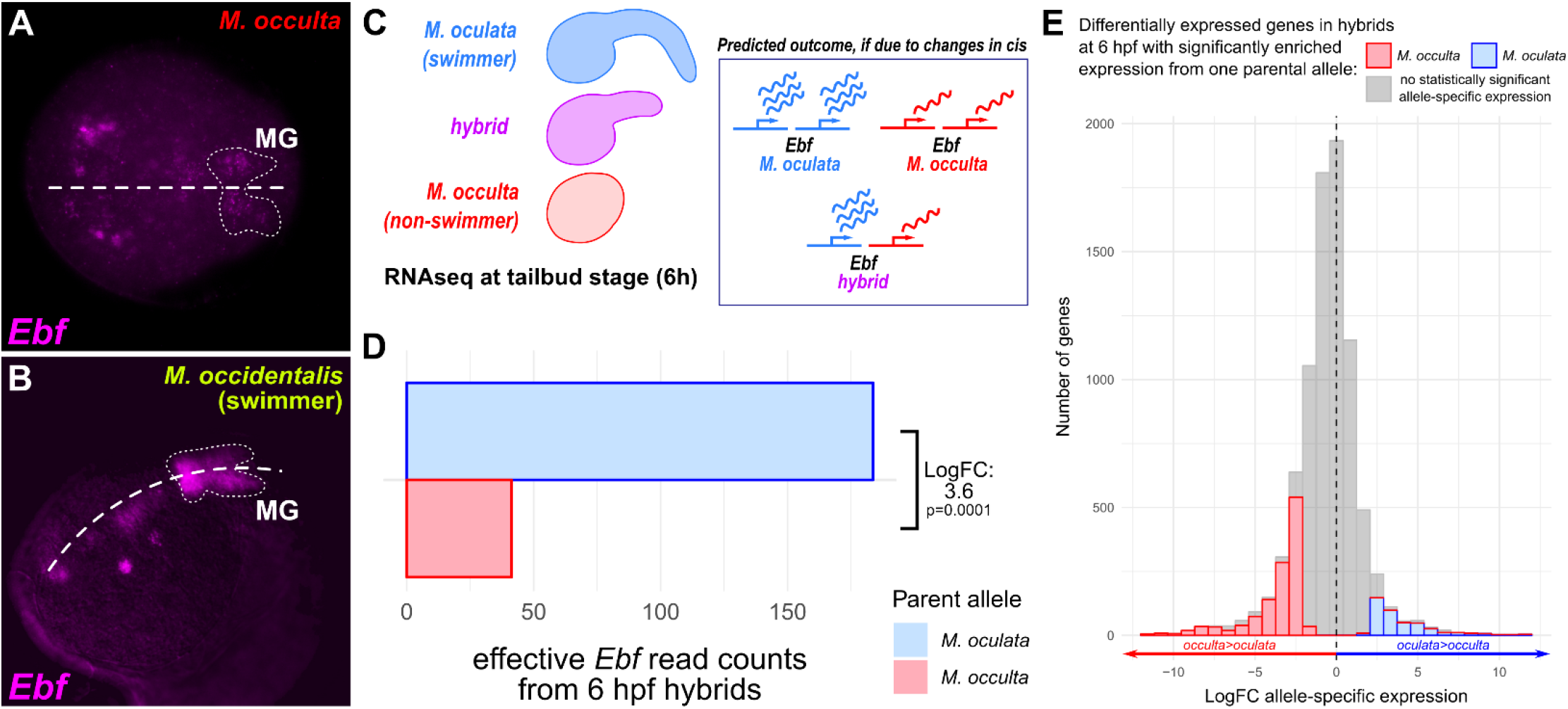
Loss of MG-specific transcriptional activity of the terminal selector gene *Ebf* in *M. occulta*. **A)** Fluorescent whole mount *in situ* hybridization (ISH) for the *Ebf* gene in tailbud-stage *M. occulta* embryo. MG outlined by dotted line, showing relatively weak expression of *Ebf*, compared to its expression in other cells in the same embryo, and to its expression in swimming species (see panel B). **B)** ISH for *Ebf* in tailbud stage embryo of the swimming species *Molgula occidentalis*, showing strong expression in the MG (dotted outline). **C)** Diagram of RNAseq analysis of hybrid embryos, produced by fertilizing eggs of non-swimmer *M. occulta* with sperm from swimmer *M. oculata* (left). If reduced transcriptional activity of *Ebf* in *M. occulta* is due to evolutionary changes in *cis*, the hybrid should show differential allele-specific *Ebf* expression that recapitulates the differential expression of *Ebf* observed between the parental species. **D)** RNAseq read counts from libraries prepared from pooled hybrid embryos at 6 hours post-fertilization (hpf), showing greater effective read count representing the *M. oculata* (swimmer) *Ebf* allele relative to the *M. occulta* (non-swimmer) allele, for a Log_2_FC of 3.6 (p = 0.0001). Embryonic dorsal midline in A and B indicated by dashed line. **E)** Distribution of genes that are differentially expressed in hybrid embryos at 6 hpf (Log_2_FC>1.5, p<0.05) according to allele-specific expression, measured and binned by Log_2_FC. Positive values indicate more abundant *M. oculata* parental allele transcripts, negative values indicate more abundant *M. occulta* parental allele transcripts. Colored bars represent those bins showing statistically significant (p<0.05) allele-specific expression, grey bars represent bins showing no such statistically significant expression. While the majority of differentially expressed genes in the hybrid at 6 hpf show no statistically significant allele-specific expression (gray), 64% of genes with statistically significant allele-specific expression show greater expression from the *M. occulta* parental allele, indicating that enrichment of *M. oculata Ebf* transcripts in hybrids cannot be explained by genome-wide bias towards *M. oculata* parental allele expression.

To compare species-specific transcriptional activity of the *Ebf* locus, we analyzed RNAseq data from embryos of *M. occulta*, the closely related swimming species *M. oculata*, and their interspecific hybrids (*M. occulta* egg fertilized with *M. oculata* sperm) (**Figure 3C**)(Fodor et al., in preparation). When we mapped RNAseq reads obtained from the interspecific hybrid to either parental *Ebf* allele, we found that transcripts of the *M. occulta Ebf* allele were significantly depleted relative to the *M. oculata* allele (Log_2_FC = −3.6, p = 0.0001)(**Figure 3D**), even though 64% of the 8306 genes significantly differentially expressed at 6 hpf actually showed allele-specific expression that was higher for the *M. occulta* parental allele relative to the *M. oculata* parental allele (**Figure 3E**) Taken together, these data indicate that evolutionary changes in *cis* have rendered the *M. occulta Ebf* gene less transcriptionally active in embryonic development.

### A *cis*-regulatory change affecting *Ebf* expression specifically in MG neurons, but not in other cells

To test for potential differences in *cis* that could explain this differential activity of *M. occulta* and *M. oculata* parental *Ebf* alleles, we constructed reporter plasmids using *cis*-regulatory sequences upstream of *Molgula Ebf* genes. Unfortunately, given our inability to electroporate *M. occulta/oculata*, we turned to *C*. *robusta*, which can be routinely electroporated in the laboratory with plasmid DNA (Zeller, 2018). When we electroporated *M. oculata Ebf* reporter constructs (*oculata.Ebf −3654/+24>XFP*) into *Ciona* embryos, we observed reporter gene expression in MG neurons, Bipolar Tail Neurons (BTNs), and some brain neurons (**Figure 4A**). Although there is no non-coding sequence conservation between *Molgula* and *Ciona*, and acute developmental system drift has rendered many *Molgula cis*-regulatory sequences inactive in *Ciona* (Stolfi et al., 2014b), in this case there was enough conservation of the underlying regulatory logic to recapitulate the expression of *Ebf* in these various *Ciona* neuronal subtypes.

**FIGURE 4 –.**
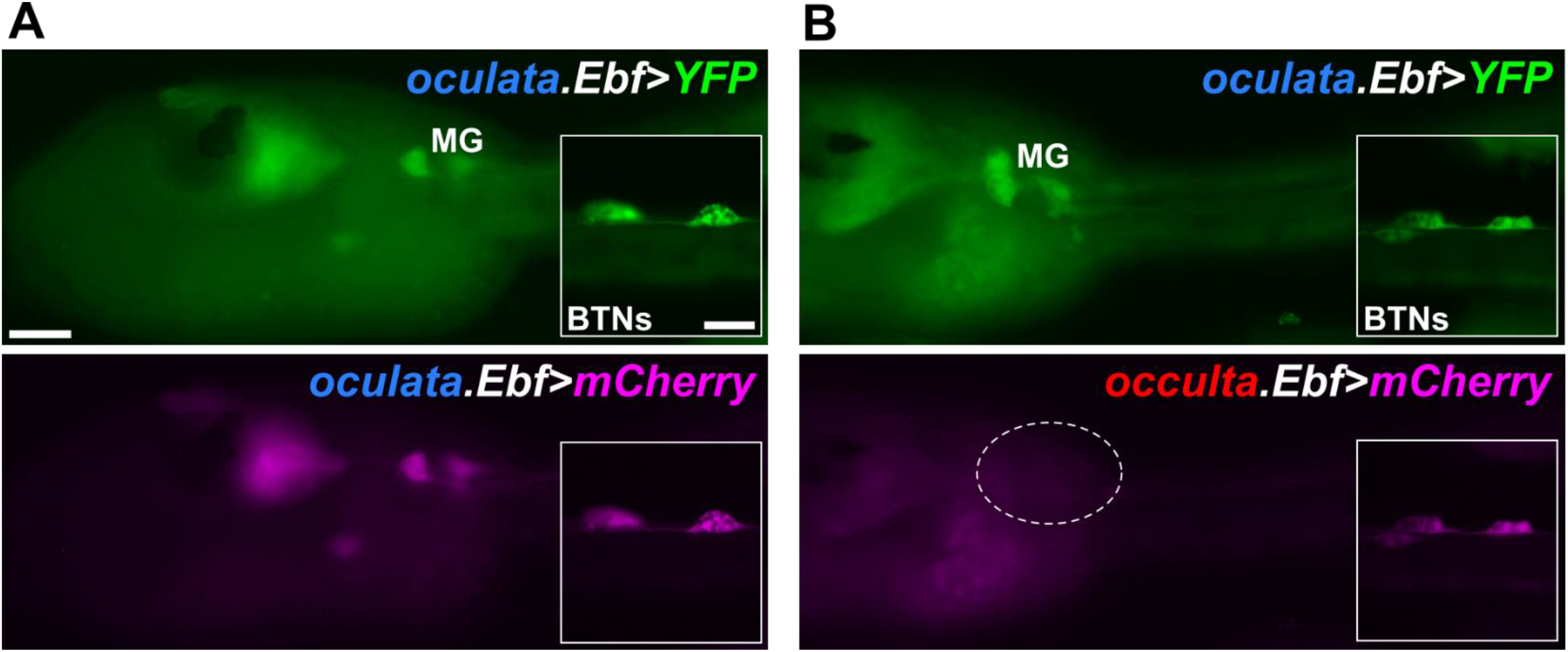
Differential activity of *Ebf* reporter plasmids in *Ciona robusta* electroporation assay. **A)** *Ciona robusta* larva electroporated with a mixture of *M. oculata Ebf>YFP* (green, top) and *M. oculata Ebf>mCherry* (magenta, bottom) plasmids, showing perfect co-expression in the motor ganglion (MG), and in other neurons including Bipolar Tail Neurons (BTNs, insets). **B)** *Ciona robusta* larva electroporated with a mixture of *M. oculata Ebf>YFP* (green, top) and *M. occulta Ebf>mCherry* (magenta, bottom) plasmids, showing absence of mCherry expression specifically in the MG (region indicated by dashed circle). In contrast, co-expression of YFP and mCherry in BTNs (insets) shows that reporter plasmid activity in other neurons is conserved. Scale bars in A: 25 μm. All panels and insets are at the same respective scales.

When we electroporated the corresponding *M. occulta* constructs (*occulta.Ebf −3659/+24>XFP*), reporter gene expression was observed in all the same neurons *except* MG neurons (**Figure 4B**) Thus, our heterologous reporter plasmid assays in *Ciona* indicated that *M. occulta-specific cis*-regulatory changes might underlie lower levels of *Ebf* expression in MG neurons, without affecting its expression in other neurons.

To identify the potential *cis*-regulatory changes underlying this difference in *Ebf* activation, we aligned 735 bp of non-coding sequence immediately 5’ of *M. oculata Ebf* start codon to the orthologous sequence in *M. occulta* (**Figure 5A**). The aligned sequences share >90% identity, consistent with the close genetic relationship between these two species. This alignment revealed an unusually divergent sequence motif ~470 bp upstream of the *Ebf* translation start codon (**Figure 5B**). This sequence was of particular interest to us because these changes were observed adjacent to a conserved E-box site that might be a binding site for Neurogenin. In *C. robusta, Ebf* is activated in the MG by the proneural bHLH factor Neurogenin (Imai et al., 2009). We identified a conserved E-box site in experimentally validated upstream *Ebf cis*-regulatory sequences isolated from *C. robusta, C. savignyi*, and *M. occidentalis* (**Supplemental File 1**), even though these sequences show poor overall conservation with the *M*.

**FIGURE 5 –.**
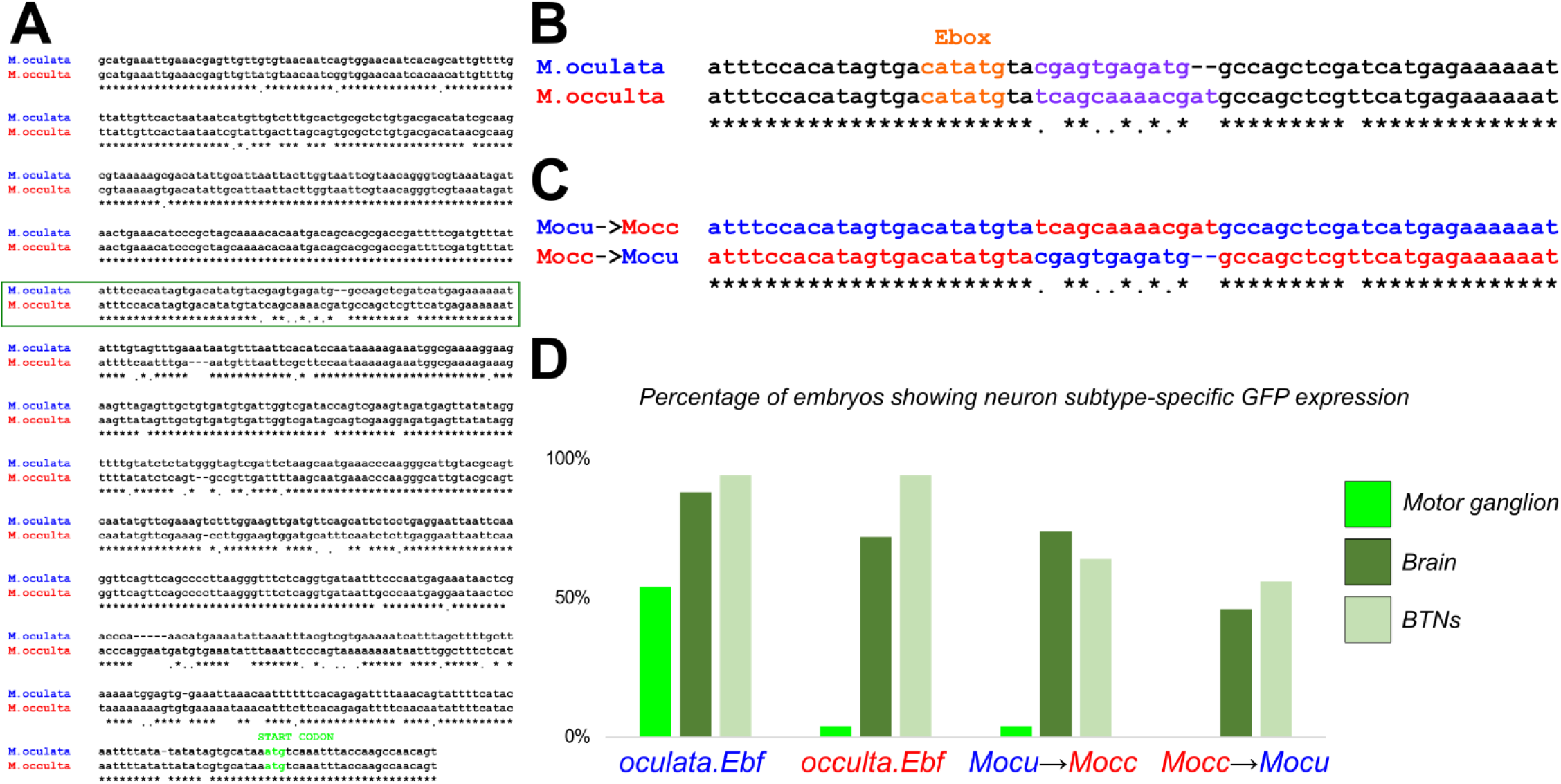
A *cis*-regulatory change underlying MG-specific loss of *Ebf* expression in *M. occulta*. **A)** Alignment of sequences 5’ upstream of *Ebf* in *M. oculata* and *M. occulta*. **B)** Sequence from boxed area in A, showing highly conserved sequences including a candidate E-box site potentially bound by Neurogenin, adjacent to a highly divergent motif (purple font). **C)** Mutated sequences in *Mocu→Mocc>GFP* and *Mocc→Mocu>GFP* constructs in which the highly divergent motif has been swapped between *M. oculata* and *M. occulta Ebf* reporter constructs. **D)** Results of scoring *Ciona robusta* larvae electroporated with the wild-type parental *Ebf* reporter plasmids and the mutated “swapped” constructs. Changing the *M. oculata* divergent motif sequence to the *M. occulta* sequence results in loss of GFP expression specifically in the motor ganglion, but not in brain or bipolar tail neurons (BTNs). The converse mutation, changing the *M. occulta* motif to resemble the *M. oculata* sequence was not sufficient to restore GFP expression in the motor ganglion. n = 50 embryos for each construct.

*oculata/occulta* sequences. The presence of an E-box within ~800 bp of the start codon in all tunicate species surveyed suggested this region might be part of a functionally conserved minimal enhancer for MG-specific *Ebf* regulation. When we compared all 3 *Molgula* sequences, we found that this E-box-adjacent motif (AGATGGC) was conserved in both *M. oculata* and the more distantly related *M. occidentalis*, but not in *M. occulta*. Because the larvae of *M. occidentalis* are also swimming, and because this *M. occidentalis* Ebf cis-regulatory sequence also drives reporter gene expression in *C. robusta* MG neurons (Stolfi et al., 2014b), we hypothesized that the E-box-adjacent motif may be crucial for MG-specific expression. Therefore, we decided to test whether the loss of the E-box-adjacent motif might explain the different transcriptional activities of swimming vs. non-swimming *Molgula Ebf* loci.

To directly test whether *M. occulta-specific* sequence changes are responsible for the loss of *M. occulta Ebf* transcription in the MG, we mutated the E-box-adjacent sequence in *oculata.Ebf −3654/+24>GFP* to match the *M. occulta* sequence (CGAGTGAGATG to TCAGCAAAACGAT)(**Figure 5C**). When we electroporated this mutated reporter plasmid (*Mocu-->Mocc>GFP*) in *Ciona*, we found that its transcriptional activity was almost completely abolished specifically in the MG, but had little effect on reporter gene expression in other neurons (**Figure 5D**). These data suggest that the mutated motif is necessary for *Ebf* expression in MG neurons in *M. oculata*. Alternatively, the *M. occulta-specific* sequence might have created a novel binding site for a MG-specific repressor, but the presence of the E-box-adjacent in *M. occidentalis* supports the former explanation. Although we attempted to “resurrect” the putative ancestral sequence by mutating the E-box-adjacent site in *occulta.Ebf −3659/+24>GFP* to resemble the *M. oculata* sequence (*Mocc-->Mocu>GFP*), this was not sufficient to restore activity, hinting at the accumulation of multiple loss-of-function (or compensatory) mutations in other parts of the *M. occulta Ebf cis*-regulatory sequence.

## Conclusions

We have investigated the molecular signatures of MG neuron specification and differentiation in a non-swimming molgulid tunicate larva, finding a surprising conservation of cell-specific expression patterns of key developmental regulators during MG development. This could indicate that at least some MG neurons might be specified and might differentiate in *M. occulta* in spite of the evolutionary loss of the larval tail and swimming behavior. Unfortunately, we were unable to investigate neuronal morphology using reporter plasmids, due to our inability to routinely transfect *M. occulta* embryos. Once this method is established, it will be interesting to revisit this question to fully document the extent of MG development in this species. Nonetheless, MG neuron function is likely seriously impaired in *M. occulta*, given the reduced levels of *Ebf* expression in *M. occulta* and of the *M. occulta Ebf* allele in interspecific hybrids. In addition to activating the expression of MG neuron-specific regulators like *Islet* (Stolfi et al., 2014a), Ebf has been shown to act as a conserved “terminal selector” that regulates the terminal differentiation of cholinergic motor neurons throughout Metazoa, including in the tunicate *C. robusta* (Kratsios et al., 2012). We therefore expect that reduced *Ebf* expression in the *M. occulta* MG would likely result in similarly reduced or abrogated expression of cholinergic effectors and other cellular machinery required for motor neuron function.

Using *cis*-regulatory mutations and heterologous reporter plasmid assays, we demonstrate specific changes to conserved *cis*-regulatory sequences upstream of *M. occulta Ebf* have affected its expression in MG neurons, but not in other neurons or cell types. If these results hold, this would represent a distinct example of tissue-specific gene expression loss in this species. Previously, it has been shown that evolutionary loss of functional larval structures in *M. occulta* correlates with loss-of-function non-coding mutations resulting in the loss of crucial proteins such as muscle actin for tail muscle activity (Kusakabe et al., 1996) or Tyrosinase for melanin pigment synthesis in the gravity-sensing otolith organ (Racioppi et al., 2017). However, these proteins are only expressed in larval structures involved in larval motility (tail muscle cells and otolith, respectively), being apparently dispensable for the development of other larval and adult tissues. In contrast, *M. occulta Ebf* is not a pseudogene and is predicted to encode a fully functional protein. This difference is likely because Ebf is a regulator that is required for the development of myriad cell types throughout the life cycle of *M. occulta*, especially in the sessile adults of this species, which are morphologically similar to adults of other species, swimming and non-swimming alike. Thus, even if *Ebf* expression is not needed in MG motor neurons (in the absence of functional larval muscles), its requirement in other tissues may have resulted instead in the evolutionary loss of tissue-specific *cis*-regulatory elements, as our data indicate. It may be that the mechanisms we show here underlying cell-specific loss of *Ebf* expression in *M. occulta* are representative of constraints acting on traits that are lost from a particular developmental stage, but that are still retained in a different stage of the same organism’s life cycle. Studying these losses and the mechanisms underlying such losses will complement the principles learned from studies of traits that are dispensable to the entire life cycle of a particular species, such as the loss of vision in cave-dwelling organisms.

Is the evolutionary loss of MG development an ongoing process in *M. occulta?* Might these processes break down further, as has happened with the loss of genes encoding melanogenesis or muscle structural proteins in this same species? Or are there special constraints that have maintained the neuron subtype-specific marker gene expression we see in the MG? Has MG neuron subtype-specific gene expression persisted because the same regulatory logic is required for expression in adult neurons or other tissues? Alternatively, we know that some MG precursors remain undifferentiated in the larva but can contribute to the adult nervous system (Horie et al., 2011). Perhaps the cell-specific transcription factor expression we observe in the *M. occulta* MG reflects shared mechanisms of patterning both differentiated larval neurons and undifferentiated adult neural precursors. Another intriguing possibility is that the tunicate MG is required for other larval behaviors beyond swimming. For instance, perhaps MG neurons are required for triggering developmental processes during the metamorphosis of the larva into a juvenile. In this scenario, specific MG neuron subtypes might still be necessary for the proper metamorphosis of *M. occulta* larvae, even if Ebf-dependent activation of motor neuron effectors in the MG is not required and might have even been selected against. A deeper understanding of the mechanisms underlying both larval and adult neurogenesis in tunicates, and the role of neural activity in the transition to the adult phase, will be needed to answer these and other questions related to the loss of larval-specific structures in non-swimming species.

## Acknowledgments

We dedicate this manuscript to the memory of Alexander “Zander” Fodor. We thank C. Titus Brown for his encouragement and continued support. We thank Stéphane Hourdez, Sophie Booker, and Xavier Bailly of the Station Biologique for help in carrying out experiments at Roscoff. We thank Susanne Gibboney for assistance with cloning *Molgula Ebf* reporter plasmids. This work was supported by NIH award R00 HD084814 to AS, by start-up package from the College or Arts and Science at New York University and NIH award R01 GM096032 to LC, by France BioImaging infrastructure ANR-10-INBS-04 to NP, and by a University of Washington Royalty Research Fund award (A118261) to BJS. The collaborative MoEvoDevo network is supported by ASSEMBLE (Association of European Marine Biological Laboratories) MARINE. CR was supported by a long-term fellowship ALTF 1608-2014 from EMBO and by a travel grant from the Boehringer Ingelheim Fonds. This material is also based in part upon collaborative work by BJS and EKL and C. Titus Brown, supported by the National Science Foundation under Cooperative Agreement No. DBI-0939454 BEACON, A Center for the Study of Evolution in Action. Any opinions, findings, and conclusions or recommendations expressed in this material are those of the author(s) and do not necessarily reflect the views of the National Science Foundation.

