## Supplemental File 1 for "A cis-regulatory change underlying the motor neuron-specific loss of terminal selector gene expression in immotile tunicate larvae"

**SUPPLEMENTAL FILE 1 – Lowe et al. 2019**

***In situ* hybridization probe template sequences (*M. occulta*)**

>Mo.occu.Celf3/4/5(ETR-1, from genomic DNA)

ATCCAATGCAAGTTAAGCCTGCGGACACCGTCAACAAAGGAGGCAAGAGAAATGCTAGTTCAATAAACGAATGTTAAAAATAATACGATTATTTTAGAGGATAGAAAATTGTTTATTGGAATGTTGGGAAAACGACAGACGGAAGAAGATGTTAAAAAACTCTTCGAACCATTTGGGAAGATCGAAGAATGCACGATATTAAGGTGAGAATTTAATCGTGTCATTAATTATTAAGAAGTAAAAATAAATTTAAATTATTTATATTGATTGCAAATGCCACATGGCTAGCGACACGGCCCACTTACACATATTATTTATATATACAGGGTGCATCATCAAGTCTTGTTGTAAAGTTAGCGGACACAGACAAAGAACGAGCAGTGAGGAAGATGCAACAAATGGCAAACAACTATGGATTAGTCAGCCCTGTGGCTTTACAACTTGGAACTTATCCAGCACATTCCCTCGTGGCACCAGGAGTCATGCCAAGTGCAGGTTGGTCACCTATAGCGACAGGAACTTTACCAACTGCTCAATTAGGACAAATGGCTAATGGCCTCAATGGCCAAACTCCAGTTACTCAGCAAAATGGTATTTGATTTAATAAGTCAATCTTTTGTTATTTATGCGAGTTTATGTACAGGTCCGACTACTCCAGGAATACCGAGCACACCACAAAGTCCTGTCGCATCGATCACAGCGTTGAACTTGGTTCCACCTCCAGTTGCTTCCCAGACAGGTTTATCATCCGGAATTAGCCATCAAGATCTGTACTCGATTCCGGCTTATCCTGGTAAGGATATAGTTCTCCTACAGTATGAGTTGCTATTTTACCATATAATCAAATTATGAGAAAATTTCATTAGAAAATGAAAATCCAATAAATTGAATGGAGGTTCATTTATTATTTATGTTTCAGCACAAACACCGCCAGCCGTTGACATGCTTCAACACCCATCATATGCTCAGCATCAACCATATACAGGTGTGTTTCTATGGCTTTAAACAATAGTGAGATAGACCTACTTTGGAGCAGTGTGCTGCTCGACAATTTGATATGGTAAACTACAATTTTAGTTGGAAAGTGCAAGTGAATTATATATGGTATAGGGTTTTGTTTTTCGCATTATCTTTCACATTCATTATTAATGATGCGCATACGCAAGCAAACGGATGTATGATTTAATGGTTATTATTTCAGTGGTGTACGTCCCATCACAGCCATATGGGGCCGGTCAGCTTGCACAAGTTGCCACAACAGGACTAACAGCCGGTGCCACTCAACTTGGACCTGCCCTAGCAACACCGCAACCAGCCACTTTAATAAACACAAGTCCCACTGCACCACAGAAAGAAGGTGACAATAATTACATAATATCAATTATTTATTTTGAAAATTTTCTGTGAAATCTAAATTAAATTTGTTCCTTTATCACTTTTTTTTTTTTAATTATGTAAATTGTGTTCCAGGGCCAGAAGGTTGCAATCTGTTTATTTATCATCTTCCGCAAGAGTTTACTGATGCTGACCTTGCCAATGTTTTCCAACCATTTGGTTCGGTTATTTCGGCTAAAGTTTTCATCGATCGAGCAACCAACCAAAGTAAATGTTTCGGTAAGCAAACTTAAATGAGTGGCAATAATGAATACTCACTGCCATGATGATGTATGGATCAAAGTTGTTGTTTGTTCAGCTTAAAGCACAACTTAACATCGGCGATAATTATAGAAACTTCCTAAGTGCATAATTTGGATATTTTTATTGCATTATGTTTATCCAATTTGTAATGGATACTCAACTGAAAGACAATTTGTCACAATGACACGGTTTAATACTCTCTTATATTTTCAGGCTTTGTTAGCTACGACAACCCACTGAGTGCACAAA

>Mo.occu.Dmbx

ACTCGGATTTGTTCTACAAGCAAGGTCAAGGGTTCCCGACTGTGGGAGGATTCCCACTGGATGTTGCGTGGCTGGGAAGGCATCAAGCATTGGTGAATAACCCAGGACTTTATCAATTCCACGACGCTTTGGCATTGGGAAAATTGGCAGACGTTATGTTGGAATCGCGTTACGGTTTACGTTCGCACAAACAACGTAGAAGTCGGACTGCTTTTACATCGGACCAACTTAAGGCGTTGGAAAATACATTTGAAGAAACCCAATATCCAGATGTGATAACTCGGGAACGTTTGGCGATTTACACAAATTTACCAGAAGCGAGAGTGCAGGTGTGGTTTAAGAACCGAAGAGCAAAATTCAGAAAACAACAAAAGTTGTTGAAGTCAAACAACAAAGGAAAAGATGTAAAGAATGATAGCGACGTTGGTGACGACAAATCAACCTCAAATGATAAATTAATCCAAATTAAAACCGATATTTTACCTAATCCGTCTACGACCGACGAGATCGCGACACCTCAATCGAATGAACAAAGCGAAAAGGCAGATTTAGGAGACTCTGACGTCACAACTGTTAACACAATGTCTCATCTCCGACACCCGTTTTCATATCCGATGTTGCCATGGTTATCATACGCACA

>Mo.occu.Ebf (from genomic DNA)

ATTCGAGTTCAAACTCCACCACGACATGTACCGGGCGTAGTTGAAGTGACACTATCGTACAAAAACAAACAATTCTGCAAGGGAAGTCCTGGTCGATTTGTTTATACAGGTTTGTTAATTCATTTGTGTGAGGTGAGGAGAGTTAAATTTAAAATGACACTTTAGCCTTGAATGAACCGACGATCGACTACGGTTTTCAAAGATTACTGAAAGCTATTCCTCGACACCCTGGAGACCCAGAACGACTTCCAAAGGTAAATTATTTCAATTAACTACCGCAAGTGACTTCCCGCGTAAATTAATCATGCAATTAATAATAAAATTTCCGTACTGGATTTAGTGCTTTCCATCGGATGAATCATTCATGCGTATCATCAACAAAATATATATTTAGGAAATTGTTTTGAAGCGGGCAGCTGACGTCATGGAAGCGGTGATGACGCGCCCATATAACCAAGTACCAGCACCTCCCCCTGCCCCCATGCACAATACATTCAATGGATCATCACCGGCAATGATGGCGGCAGGAGTCAACACGTACAATCACAGTGTGCAGATGGCACCTTCACAGTACACCATCACTACACCAGATCGTCTGGATTCAGCCAACGGTAGCGACTCCGGT

>Mo.occu.Islet

AACAAAATATACCACATCGACTGTTTTAGGTGTGTTGCGTGCAGTCGACAGTTGATTCCTGGTGATGAATTTGCTTTGAGAGATGAAGAACTTTTCTGCAAAGCTGATCATGATGTTGTGGAGAGAGGAGACGTGTTAGCGATGCCTCCTGGAGATATGTCGTCGTTGACGGGATTGTCACCTCCTATGAGCATACCTAACGGCGGGGTTCTAAGCCCAGGGATCGGCGGGATTCCCGGTGGAATGTGCGGCACGAACAGCTCCGGAAACGGCTCATCCAGTTCCGGAAGTCGTCGGTCTCATAAGGATCCCAAAACGACGAGAGTGCGAACAGTATTAAACGAAAAACAATTGCACACACTTCGGACGTGCTATGCAGCGAATCCGAGACCCGATGCACTCATGAAGGAACAGCTTGTCGAAATGACGAATCTATCGCCTCGCGTCATCCGGGTTTGGTTTCAAAATAAAAGATGCAAAGACAAGAAGCGTGCAATTGCTATGAAGCAAATACAGGAACAGAACCAGCAACATCAGCAAGGAAATGAAAAGAACCATGGTGGACAGAAAGGTCTCTCTGGTATGAACGGAGTTGCTATGGTTGCGAGTGAACCGGTGCGAAATGATAACCCTGTCAACAACGCCAAGCCATTGGAAGTAAGAAATTATCAACAACCAGCGTGGAAAGCACTCAGCGACTTCGCTCTTCAAAGCGAGATAGAACAACCTGCATTTCAACAACTGGT

>Mo.occu.Mnx (from genomic DNA)

TTAGCGCCCAAAGTCATAATGGAGTAATGCATAGAAGCATGAATGAATATCTAACAAGATCAGCCGATTTGTCTCCTCCTGAAGAAGACAAAGTTTGTCGATCATACCACTCTTCTGAAGATGATGTTCAAAAAAAGACGGTAAATCGATCAGCTTCAACATCTTCATGGTCATCAGGATCCTCTCTCACTTCACCATCTCGATATCCAAACGACCACGTTCAAAACCACAGTGGAAGCAACGGCTCATCATCGGATGCACAGCTGGAACACGGAAAAATATCGCCAAGAGGAGCGCAATTCAGAATCGAATCATTACTGTCGATGTCACCAAAAAATTCGGATATGAAGAATGAATTAAACAATGACAATAAACTGGATTGTCGTTGGTACGACTTTGTGAAGAACAATGGAAAAATCAAACAAAATGGCGGTTCGAGACGCCAATCACAAGGCAGCGATATATCCTCAAAGGTTTCGCCAACGTCAAGTCCTTATATAGACGTTGTAAATGACGATGGCAACGAACGTCGCGGGTCTTTGACGGATATTTATCCAGGAATCCCCAAACCCAACAGTTCGGCTTTCCAATCCACCGAAGCTTCATGTGGAATGCCTCCTTTTGCAATGTACCCAAACTTCCCTGGAGTTCACTTTAATCAGTTATCGTTTGCCGCGGCATTGGCAGCTTCACACATGGAAAGAAATGCGCAAATCTCAGCAGAAAGCAATCAAACTGTTGCCAATTTAAGTAAAAATGAGAAAGATGCTTCCAAGCCAAGCGGCCACCCACCCATGGTTTATCCTAGTCCCTTTTCAAACTTGATCCCGTCCATTGGACAACACCCATTTCTTGGCCAATATGGGTTCCCACAAGCGACAGCCGTTACATCAATGCCAGGTTACCCAGCAGGATCACATAACTTCAACCTTCCCCATTCGTCCTCCCAACATCCACAGCAAACTCTTGCCATGGAACTACTGAGGAGTGGTCGCATGTTTCGAGA

>Mo.occu.Neurog

AGCAGTTATGGAGGATGAGAAGAAGAATGAAGATTCTAAAAACAAGAAAAAGCGAAAGCGCGAAAGAAGTCCGGAATCAACGGTTATCCTGAAGAAGATTCGTCGTGGAAAGGCAAATGACAGAGAACGAAATCGCATGCATGGACTAAACGATGCGTTAGAAAATTTAAGGTGAGATTTGCAAATTTTTTTTTTGTTTTTTCGTTATTTTAATGCTATGTTTTTAGACGAGTTTTACCGACATATCCCGACGAAACAAAACTGACGAAGATCGAAACTTTGCGTTTTGCATACAATTACATTTGGTGTTTGAGTGAAATGATTAAAGGCAGTGAAAACGATCCAAACGCAGCAGGTAAAAATCTCATTAATTATCATTAGTAATTAACCGCTCTTGGATACCATGTTCTGTTATTATAACGTCTGCTTACACAGATTTGGCACAAGCGGCCTTCAACATGCAGCAAATGCCATCGTTCAGTGAAAACATCGTCTCCTCGACAACTTTGCATCCCATCGACGCCAATTCGCCTCCGATGATGCCGTACAATGAGTACCAACAACAGACCACATTCTCCGCCGACCAGCGACCGACTTACGGAAGTTTCGGAGGTGATTCTCGAATACACGGAGCTCATCTGCAGCAGTCATTGGTTGCAAATGAAATGGCAAATTGTCTTGATGAAATAGAAGATGTTTCACAACCATTCATCACACCATACGATCAACGACATACAAGTAGTGACGTCGGACAGCAAGGGATGGTTTATGGCAACGAAATGGCAACGTATCAATCCGCTACAACAATTAACAACAATAATAGCAACAACAATAACAACAACAACACCACTGCATTCACAAATCTAAATTCAATTGGATACCGTCGAAATGCAAAGGAAATTTCACTGCAAGGTTTAAATCAAGTGTCCCA

>Mo.occu.Nk6 (from genomic DNA)

TAACGCTCACAGGATAACATGGATAGCTCAGCAGCACAGGGGGCGTTTTTGTTCAATAATAATCACGTTGCCGCGGCCATGTCGGCCCTACAGGCAGCATCGTCCGGAATTAATGAAACCAGATTGCAATCTCCAAGTTTTCATTCCTATTCCCCGTTTCAGAACGGTGCAGTTTTTCCAAGCATTCAAAAATCCCTCAACAGTCCAACGTCAGCGACGCCGTTCGGAATCAACGATATTTTGAACCGCCCAAACACTTCCACGGAGTCGCTCCAACCAAGGACGAATTCACCTCCAGGACCATCTGGTAACGTTGCTGGTTACTTCTCCAACGGCCACAGCCACGTCACTACACCAGGAGCAGCCATAGCAGCGGCAGCAGCAATGTATCTCGGTGCAAGTTCAATAGGAGCATCCAACAGCGTATCTGGAGGAATGCTGACACAGAATCCCGCTTCAAGATACGCCAAACCACTCGCAGAGCTTCCAGGAAGAGCTCCAATTTACTGGCCCGGAGTTCTCCAGAAAGATTGGCAGGAAAAGTTCTCATGTCAAGGTAACGCGTTGTTGTTTAATTGCTGATGCTGTGTTTAAAATGGCGTCATGATAACATGATAGTTCTTCGTATTACTGTCACAGCTGTATAACATGATGTCATCGACATCAACAAACATGTTATTATCGCATTATCATTGTTGATTTTAGGTGGACATGCTGGTTTGGCCATTGACAAATATGGTAAAAAGAAACATACTCGACCAACGTTCTCTGGACAACAAATCTTTGCACTTGAAAAAACATTTGAACAGACCAAATACTTGGCTGGACCAGAAAGAGCGAGATTGGCATAT

>Mo.occu.Onecut

ATATGACGACGGCTCCTGTCCTGACAGATTGCGTGATGGCTACCAAAGGCAATCACAAGTAAACATGTTTCCCAACCAGACATTTCTCAATAATGGAGGATTTTCAGGCGATTTAGGAAACATTCAGAACACCTCAGCTGCTGATAGCTACGCAACCCTTCAAAATGACCAACAAGACCCTGGGTTAAGTTACGCGACCCTGACCCCACTCCAATCTTTACCGATAACTTCATCTAGTGGTGATAAATTTGTCCCTGTGCCCGTCAGTTCAAACTTCCCACTTGGTAACCCTCCGGATTCCATTGATTTAAATGGGAATTATCAAAAAATGACTGGAATGGGTCAAAGTCTTCCCCCTCTTTCTAACAGCATGTTATTAAATGGTCTTCCCACTGCCACTGATAGCGTTCACGCGCCAACGACGAGCCACGTTTCACATCAATCAGACGAAATCCCATATACCGCATCAGTGATGCATCTTCCGCAATACCCTCGATCGCCTGGTAGTTTTACCGGAACGAATCCTTACGATGCCCGAGTTTTTGACGCTGTCTCCGACACTTTTACTAACCCGATGTTCCCAGGACGGACCACAGGTTTTCCAACACCTAGCATACACAACACACGCTCGCCCATCAACACTCGCGTTAACAACAGAGGACCCAGGGGATCACCAGTCAATCTTAATAATGGGAACCAACGTCAACAAAACACTGAGGAGGTTAACACAAAAGAAGTTGCGGCGAAAATTACACAAGAACTTAAACGTTACAGCATACCACAAGCAATATTTGCACAAAGAGTTTTGTGTCGAAGCCAGGGAACACTTTCGGATCTTCTCCGCAACCCAAAACCTTGGTCCAAGCTAAAGAGTGGTCGGGAAACGTTTAGAAGAATGTGGAAGTGGCTACAAGAACCGGAATTTCAACGAATGTCGGCACTTCGACTA

>Mo.occu.Vsx (from 5’RACE)

ATCACATTACAAAATGGAAGAAAATAAAGTCAAAGTAAAATCTTGGAGAGAAAGTTTTAGCATTGAGTATTTGCTGCGACCAGAGAAAAGAAATAACATAAAAGTGATGGATACAAAAATTCATGGAAAAGACCAATTGAATGCAATTTGTGCTGCGTTGCAATATTTCCAAACAATTTCTGAAAATTCGAATAAACCATCGAACATTATTTACCAGAATGAGAAGACGGTTGTTACAATTAAACAAGAAGAAAATGAAATTGAACAATTAAATCAGCACAAACAACTGGCAATAAAAGAAACGAAAAAGGAAGGTACTAAAAGACCCACATTAACCAAGCCACGACGAAACAGAGCAGCGTTCACTGAAAATCAATTTAAAATGTTGGAAGAGGCATTTAATCTTTCGCATTATCCTGATGTTGCTACCAGAGAAGAACTTTCACGGAAGAGCAACATTGATGAGACTAGGATTCAGGTTTGGTTTCAAAATCGACGTGCAAAATGGCGTAAAACATCGAATGATTGGGGACGCAGCAGC

**Proximal *Ebf cis-*regulatory sequences from *Molgula* spp.**

“Mocu” = *M. oculata* (swimming)

“Mocc” = *M. occulta* (non-swimming)

“Moxi” = *M. occidentalis* (swimming)

START codon (coordinates for each are given in bp from start codon)

>mocu.ebf-736

GCATGAAATTGAAACGAGTTGTTGTGTAACAATCAGTGGAACAATCACAGCATTGTTTTGTTATTGTTCACTAATAATCATGTTGTCTTTGCACTGCGCTCTGTGACGACATATCGCAAGCGTAAAAAGCGACATATTGCATTAATTACTTGGTAATTCGTAACAGGGTCGTAAATAGATAACTGAAACATCCCGCTAGCAAAACACAATGACAGCACGCGACCGATTTTCGATGTTTATATTTCCACATAGTGACATATGTACGAGTGAGATGGCCAGCTCGATCATGAGAAAAAATATTTGTAGTTTGAAATAATGTTTAATTCACATCCAATAAAAAGAAATGGCGAAAAGGAAGAAGTTAGAGTTGCTGTGATGTGATTGGTCGATACCAGTCGAAGTAGATGAGTTATATAGGTTTTGTATCTCTATGGGTAGTCGATTCTAAGCAATGAAACCCAAGGGCATTGTACGCAGTCAATATGTTCGAAAGTCTTTGGAAGTTGATGTTCAGCATTCTCCTGAGGAATTAATTCAAGGTTCAGTTCAGCCCCTTAAGGGTTTCTCAGGTGATAATTTCCCAATGAGAAATAACTCGACCCAAACATGAAAATATTAAATTTACGTCGTGAAAAATCATTTAGCTTTTGCTTAAAAATGGAGTGGAAATTAAACAATTTTTTCACAGAGATTTTAAACAGTATTTTCATACAATTTTATATATATAGTGCATAAATGTCAAATTTACCAAGCCAACAGT

>mocc.ebf-739

GCATGAAATTGAAACGAGTTGTTATGTAACAATCGGTGGAACAATCACAACATTGTTTTGTTATTGTTCACTAATAATCGTATTGACTTAGCAGTGCGCTCTGTGACGACATAACGCAAGCGTAAAAAGTGACATATTGCATTAATTACTTGGTAATTCGTAACAGGGTCGTAAATAGATAACTGAAACATCCCGCTAGCAAAACACAATGACAGCACGCGACCGATTTTCGATGTTTATATTTCCACATAGTGACATATGTATCAGCAAAACGATGCCAGCTCGTTCATGAGAAAAAATATTTTCAATTTGAAATGTTTAATTCGCTTCCAATAAAAAGAAATGGCGAAAAGAAAGAAGTTATAGTTGCTGTGATGTGATTGGTCGATAGCAGTCGAAGGAGATGAGTTATATAGGTTTTATATCTCAGTGCCGTTGATTTTAAGCAATGAAACCCAAGGGCATTGTACGCAGTCAATATGTTCGAAAGCCTTGGAAGTGGATGCATTTCAATCTCTTGAGGAATTAATTCAAGGTTCAGTTCAGCCCCTTAAGGGTTTCTCAGGTGATAATTGCCCAATGAGGAATAACTCCACCCAGGAATGATGTGAAATATTTAAATTCCCAGTAAAAAAAATAATTTGGCTTTCTCATTAAAAAAAAGTGTGAAAAATAAACATTTCTTCACAGAGATTTTCAACAATATTTTCATACAATTTTATATTATATCGTGCATAAATGTCAAATTTACCAAGCCAACAGT

>moxi.ebf-739

TTCATTGTTAAAAGTAAATTTAAATTTGGTTAATGTTACACGACATAACGTCAGCGAAAAAACGCCATATTGTATTAATTATTGGGTAATTCGTTTCGGGGTCGTGAAATCTTTAACGAGTAAATAGATAAATATAATGTCCCGTCGCCTGATAACAATGGACGCGTGACATTTGTTTATTGTTTATATTTACAGTAAAAACAACATATTGTAGGAAAAGATGGCAGATTTCTTCGTCTGACTCGCAGTTCTACAACGTTGTGTAAAAAGAATTAAACATTCCCTAAAGAAATACTGCTTATAATAAAACAAACAAAAATAAAAAGTCGCGAGAGAAAGTTATGAGATAGTTTTTCATTGGTCGATAGATTTATATCAGATATCTCGAGCGAGTTTTATAAACGTCCCCAACGGTACGGCTGAGTTGATCGAGGTAGGCCCATGGGCATTGCAGATGGTTACTCTTTGATTAGAACCACATCCAACTATACAATCATCTCAGGAGGAATTAAAAATAATGCTTGAAAAATATCATTCAAGTTACGCAAGATGCCCATGCTTGTCCCTCGAGTTCATTGTGACAAAAAACTTGCCGTTGGAAAATGCAGAAAAATTATGTCAGAAAAAATTTTTAAATTAAAGTCAAGATTTAGGTTTTAAAGTTTGCAATAAATACATTTTTGTTTTACAGTTGTTAATCTAATTTAAAGCAATATTTCGGGATAATTAGGTCGATATATGGCGTCTCTT

**Alignment of *Molgula Ebf cis*-regulatory sequences**

E-box

E-box-adjacent motif

Ultraconserved motif

```
moxi.ebf-739    ------------------------------------------------------------
mocu.ebf-736    gcatgaaattgaaacgagttgttgtgtaacaatcagtggaacaatcacagcattgttttg
mocc.ebf-739    gcatgaaattgaaacgagttgttatgtaacaatcggtggaacaatcacaacattgttttg


moxi.ebf-739    ---ttcattgttaaaagtaaatttaaatttggttaatgttac---acgacataacgtcag
mocu.ebf-736    ttattgttcactaataatcatgttgtctttgcactgcgctctgtgacgacatatcgcaag
mocc.ebf-739    ttattgttcactaataatcgtattgacttagcagtgcgctctgtgacgacataacgcaag
                   **  *...*** *.* .  **.  ** *    ..*.* .   ******** **. **

moxi.ebf-739    cg-aaaaaacgccatattgtattaattattgggtaattcgtttcggggtcgtgaaatctt
mocu.ebf-736    cgtaaaaagcgacatattgcattaattacttggtaattcgtaacagggtc----------
mocc.ebf-739    cgtaaaaagtgacatattgcattaattacttggtaattcgtaacagggtc----------
                ** *****..* *******.********.* **********  *.*****

moxi.ebf-739    taacgagtaaatagataaatataatgtcccgtcgcctgataacaatg--gacgcgtgaca
mocu.ebf-736    ------gtaaatagataactgaaacatcccgctagcaaaacacaatgacagcacgcgacc
mocc.ebf-739    ------gtaaatagataactgaaacatcccgctagcaaaacacaatgacagcacgcgacc
                      ************ *. **..*****... * .*  ******  ..*.**.***

moxi.ebf-739    tttgtttattgtttatatttacagtaaaaacaacatattgtaggaaaagatggcagattt
mocu.ebf-736    gattttcgatgtttatatttccacatagtgacatatgtacgagtg--agatggccagctc
mocc.ebf-739    gattttcgatgtttatatttccacatagtgacatatgtatcagcaaaacgatgccagctc
                  * **.. *********** **   *. .  *.**.*   ** . .* .  ** ...*.

moxi.ebf-739    cttcgtctgactcgcagttctacaacgttgtgtaaaaagaattaaacattccctaaagaa
mocu.ebf-736    gatcatgaga-----------------------aaaaatatttgtagttt-------gaa
mocc.ebf-739    gttcatgaga-----------------------aaaaatattttcaattt-------ga-
                  **.*  **                       ***** * **  *  **       **

moxi.ebf-739    atactgcttataataaaacaaacaaaaataaaaagtcgcgagagaaagttatgagatagt
mocu.ebf-736    ataatgtttaattcacatccaataaaaagaaatggcgaaaaggaagaagttagag-----
mocc.ebf-739    --aatgtttaattcgcttccaataaaaagaaatggcgaaaagaaagaagttatag-----
                  * **.***   ..   * **.***** *** .*. . .**..*.*. *   **

moxi.ebf-739    ttttcattggtcgatagatttatatcagatatctcgagcgagttttataaa---cgtccc
mocu.ebf-736    ttgctgtgatgtgattggtcgataccagtcgaagtagatgagttatataggttttgtatc
mocc.ebf-739    ttgctgtgatgtgattggtcgatagcagtcgaaggagatgagttatataggttttatatc
                ** ...* .  .*** *.*. *** *** ..    ....***** ****..   ..* .*

moxi.ebf-739    caacggtacggctgagttgatcgaggtaggcccatgggcattgcagatggtta--ctctt
mocu.ebf-736    tctatgggtagtcgattctaagcaatgaaacccaagggcattgtacgcagtcaatatgtt
mocc.ebf-739    tcagt--gccgttgattttaagcaatgaaacccaagggcattgtacgcagtcaatatgtt
                .      .. *..** *. *   *.  *..**** ********.* ...**.*   * **

moxi.ebf-739    tgattagaaccacatccaactatacaatcatctcaggaggaattaaaaataatgcttgaa
mocu.ebf-736    cgaaagtctttggaagttgatgttcagcattctcctgaggaattaa--------------
mocc.ebf-739    cgaaag-ccttggaagtggatgcatttcaatctcttgaggaattaa--------------
                .**  .   ... *  . . *.. .  .  ****  **********

moxi.ebf-739    aaatatcattcaagttacgcaagatgcccatgcttgtccctcgagttcattgtgacaaaa
mocu.ebf-736    --------ttcaag-----------gttcagttcagccccttaagggtttctcaggtgat
mocc.ebf-739    --------ttcaag-----------gttcagttcagccccttaagggtttctcaggtgat
                        ******           *..**  .. *.****..**  . *. ...  .*

moxi.ebf-739    aacttgccgttggaaaatg--------cagaaaaattatgtcagaaaaaatttttaaatt
mocu.ebf-736    aatttcccaatgagaaataactcgaccca-----aacatgaaaatattaaatttacgtcg
mocc.ebf-739    aattgcccaatgaggaataactccacccaggaatgatgtgaaatatttaaattcccagta
                **.*  **. **...***.        **     . ..**  *     ** **.  . .

moxi.ebf-739    aaagtcaagatttaggtttt----------aaagtttgcaataaatacatttttgtttta
mocu.ebf-736    tgaaaaatcatttagcttttgcttaaaaatggagtg-gaaattaaacaattt---tttca
mocc.ebf-739    aaaaaaataatttggctttctcattaaaaaaaagtgtgaaaaataaacattt---cttca
                 .*.  *  ****.* ***.          ..***  * **   *   ****   .**.*

moxi.ebf-739    cagttgttaatctaatttaaagcaatatttcgggataattaggtc----------gatat
mocu.ebf-736    cag---------agattttaaacagtattttcatacaattttata-tatatagtgcataa
mocc.ebf-739    cag---------agattttcaacaatattttcatacaattttatattatatcgtgcataa
                ***          .****  *.**.*****. . *.****  .*            ***

moxi.ebf-739    atggcgtctct-------------t
mocu.ebf-736    atgtcaaatttaccaagccaacagt
mocc.ebf-739    atgtcaaatttaccaagccaacagt
                *** *.  *.*             *
```

**Proximal *Ebf cis*-regulatory sequences from *Ciona* spp.**

START codon

>Ciona robusta

TTTAAGCCGACAAAGGCCACAAAACTGCCTAAAATTTATAAAAAAAATGTAAAATTATTTTGTTTTTTTTAAACTACAGTTATCACCTTTAAAACAAACAAATTAGCAAACGTTGTAATTACTTCACAACTTTCTTGCGACGCTAAAAGGCGGCGAATTTTATTGCTATTGTGACGTCACAAGCGCTCTCGTCACGCCCGGATACGATTAGAACAACGAAGGATTGTTTGTTTTTAATTATTTCTCTGTTTAATCATTTGATTTAGCGCGGCACAAATTTTGTTTTATATAAAATGTATCCATTTTATCTCTGCGCGTTTTTGTACTATTTTTTGAAAAATGTTTGTTAACCTTTTGAAAATCGCGAAACCAACGAAATTATTTCCAAAGGCTGTACAATTCTTTTTCGTTAGGTTACGTGTTTAAGTATAGGCCCAAGTTTTAAATGGCGGACAGAGTTTCGTATTTTGATATTTGAATTTTTTTGAAATCTTGAAAAAAAATAATTGTACGTTTTACATAGAATAACTAACCAAAATATCTGAAGCAAAAATTACGTACAATTTTTTAAAATGTAATTTACTTTCTAGCTTTTAATTTTTGTGCTTTTCTCAATATGTGTGCCATATTTTAAAAACGTAAATTTGCCTGTTGTTAAGCGGAGGAAAAAGTAATCTGCGTGAATGCGAAAAACACGATTTTGGAATCAGCGCCGGCAATGGTGTTTGTAAATAGGGGTGGCATACGCGTTTCGTAGCGAAAGAGAGAATGAGGCGAAAGTCGACAGATGCACGCTCCGATTTATGAGACAGGAACCAGTCGCGAGGGCACGGAGGAAAAAGAACCTTACTCCAAGACATGCGCCGCCTTTTTTCTTTCTGTCCTAGTCAGGAATACTAGAGTATAGAAGGCCACGCGTCGTGGAGTTTAAAACCAGCAAGCCAGTGTCTCAACGGACACATCAATACAGAGACTTCTCTCAGTGGACAACTCGGACGATTCGCCACTAACTTGGTGGATTCGTCCCGGACGACCCATGGGCCGGGTCCCAGCGCGCTAGTTGGCCACCATACAGTGTAGAATCAGCTAGATCGTCTCTGCGGATTTCGCAAATAGATTGAGTTGGAGATAGCTTCCCGACCGGGATTTCGACTAATTTGCAATGTTAGTTATTAATCAAGGTGACAGTCAGGAGTTAAAGTTAATTTACCTTTTGAAAGGGCAAAAAGATTTTCGAAGTAAATTGATTCCGTTAATTGTAACTCTTAAACGCAAACCGATAAACGACGCCATTTTGCTTTTCATTGAGAAACTAACCATTTAGGCATTCTATAATTAAAATTAAATGTTTTTAAAATCTGTAACTATCTGTAATAAACTAGAATTATTTATTTCAGTTTAAATTTTTATTTAAAACAATAATTTTATTATTTCCTTTATTCATCCTAATGATATTGGTATAGAGAAAACGATGTTTTTATTTCCATAAAATTTTATTTCAGAAAAGTCATTTTGTTCAATTTAAAAAATTCCTTTATTCTAAAAAAATGCCCAAAACAAGTTCTTTATTTCTTAAAAATTATCCTAAATAAAAAAATCCACGTTTTAATAAAATGTATAAAATTGAAAACTATAAAAAGATGACTTATTTTTTTACCCTAACGTGATTTTTCACCAGATACCTTAAGTGTTATTTTATTTGTAAGTAATATCCAAATGGCAACAATCGCG

>Ciona savignyi

CAATAAATTTAAATAAACAACATACATTCCTTTATTTCTACAGAAATATGATAAAATTTAGACACGATAATTTTTTTACACTTTAGTGTAACTATAAACTTGGTTAAACAAAGTATCAACCGAAAAAAATATTTTTTTTATATTGCGGTAATTATAATTGGCCGCGGTTATCATTAGTTCCAAATCACATATTTTGATTTTGTCAGCTCTAAGTTTAAAAAAACAACAAATAAAAGTGTGGTAGGGGTAGTAAAATTAAAGACCGCGATATGTTTTAGCTAGTTGTCACCCGTGGAACAAATTACAGGTTTATTTAACACCAGATCATCACTTGTATTTTTATTGCTATTGTTTCAAATCATTTCGCCCGTCACGCCCGGATACTATTAACACAATCATTAATTGTTTGCTGTAACTATTACGCGCGCCAATCTTGTTTACATTTGAAAGAGAGAGCCGCATTTTCGTGTCAAAGACACGCAATTCAATTTTAATTGCCACCTACAACAACAACATGTGTTAAACAAATAATAATATTTATGGCTGTTCGTTTTAAACTGATTCTATACGATCTATAAATTATCTCTGGAATATAACGAGCAGCTTCAAAATGGCGCCGTCCCTTCAACATGTTCTCTGATTCTACAAATATATTTTTAAGGAAATATTATAGAAATAATAACTTTCACCTCCGACGTATTTTACACGCGCCGACTATTTGTTTTACTTTATTGTCATAAAATAAGGTTTTTGCTTTATTATTACATTACTGTTAAAAATAGGAGATAGGAAATTGTGTGTGATATTCAGGCAGATCGTATTGGAAACTACGAAGTAATTTGTAAATTGAGGGCAGCATACGCTGTCCGGGAAGCTAACGAGAGAAGCAAAAGTCGACAGATGCGAAAGCTCCGATTTACGAGATACCAGGCAATCTGGGGGAGAAACGCAACTTGTTCTCGTCTTTTTTCCTCAATCCGAACGGGAATACTACAGTATAGAAGGCACGCGCCTCGCTGAGTTTAAAACCAGCGAGCCCAGTGTCTCAACGGACACATCAATACCCAGCGTTCTCTCAGTGGACAGTCAGGACGAACTTTCAGAACTCGTCTCGCTCGACCCAAGGCGACGGGTCCCAGCGCGCTAGTTTACTGCCTAATAGCGACTGCAACAGTGACTTGTCTCTGTGGGCTTCGGAAATAGATTGACTTTGAGTTTTTTTGACTTTCGTTTAATTTTCATTACGTAGTTATAATTAAGACGATCGAAGCATTGCGATTAATTTCTATGAATGACAAACGTTGTCTGGAATAATCTGTTTCGTAAATAACAGGAAAATATATATTTTTTAATATCCTAGTCATATACGTTATTACAAGTGGTAGTTTTCTCCGTTAAATTGTTTATTTAAAAAGACGTTGCTTTAATACAGTAGGCTACTAACTAAATACTTATTCAAAAGCTATTTTTACATTAATTATTTTGGTATGTTAGTTACCATTATATCGAAAACATTTCAACAGATTTTGTTTGAATTAATTATTTGTGAAGTAAAATATGGATGGACCTCAACTTCCAGGCTCAGCGGTACGTGGTTGGATGCAAACTAGTCTGATTGAACCAATGGCAAG

**Alignment of *Ciona Ebf cis*-regulatory sequences**

```
E-box

Robusta         -------tttaagccgacaa----------------------------------------
Savignyi        caataaatttaaataaacaacatacattcctttatttctacagaaatatgataaaattta
                       *****.. .****

Robusta         ------------------------------------------------------------
Savignyi        gacacgataatttttttacactttagtgtaactataaacttggttaaacaaagtatcaac


Robusta         ------------------------------------------------------------
Savignyi        cgaaaaaaatattttttttatattgcggtaattataattggccgcggttatcattagttc


Robusta         -aggccacaaa-----------actgcctaaaatttataaaaaaaatgtaaaattatttt
Savignyi        caaatcacatattttgattttgtcagctctaagtttaaaaaaacaacaaataaaagtgtg
                 *...**** *            * **.. **.**** ***** **.. * **  .* *

Robusta         gt---tttttttaaactacagttatcacctttaaaacaaacaaattagcaaacgttgtaa
Savignyi        gtaggggtagtaaaattaaag--accgcgatatgttttagctagttgtcacccgtggaac
                **     *  * ***.** **  *.*.*  *  .  . *.* *.**. **  *** * *

Robusta         ttacttcacaactttcttgcgacgctaaaaggcggcgaattttattgctattgtgacgtc
Savignyi        aaattacaggtttatttaacaccagatcatcacttgtatttttattgctattgtttca-a
                  *.* ** . .* *.* .*. *.    *  .*    * ***************  *.

Robusta         acaagcgctctcgtcacgcccggatacgattagaacaacgaaggattgtttgtttttaat
Savignyi        atcatttcgcccgtcacgcccggatactattaacacaatcattaattgtttg-ctgtaac
                *. * . * *.**************** ****. ****. *  .******** .* ***.

Robusta         tatttctctgtttaatc--atttgatttagcgcggcacaaattttgttttatataaaatg
Savignyi        tattacgcgcgccaatcttgtttacatttgaaagagagagccgcattttcgtgtcaaaga
                **** * *   ..****  .***.  ** * . *. * *. . .  ***..*.* *** .

Robusta         tatccattttatctctgcgcgttttt----gtactattttttgaaaaatgtttgttaacc
Savignyi        cacgcaattcaattttaattgccacctacaacaacaacatgtgttaaacaaataataata
                .*. ** **.* .*.*.  .*.. ..    ..* .* . * **  ***..  *. ***.

Robusta         ttttgaaaatcgcgaaaccaacgaaattatttccaaaggctgtacaattctttttcgtta
Savignyi        ttt-----atggctgttcgttttaaactgattctatacgatctataaattatctctggaa
                ***     ** ** .  *   . ***.*. ***.* * * * **.** *. *.*..*  *

Robusta         ggttacgtgtttaagtataggcccaagttttaaatggcggacagagtttcgtattttgat
Savignyi        tataacgag--------------cagcttcaaaatggcg---------------ccgtcc
                 .* *** *              **. **. ********               ..   .

Robusta         atttgaatttttttgaaatcttgaaaaaaaataattgtacgttttacatagaataactaa
Savignyi        cttcaacatgttctctgattctacaaatatatttttaaggaaatattatagaaataataa
                 **..*  * **.*  .**..*. *** * **  **. . .  *  .******  * ***

Robusta         ccaaaatatctgaagcaaaaattacgtacaattttttaaaatgtaatttactttctagct
Savignyi        ctttcacctccgacgtattttacacgcgccgact------atttgttttactttattgtc
                *.   *. **.** *.*     .***..* . .*      ** *. ******** * *..

Robusta         tttaatttttgtgcttttctcaatatgtgtgccatattttaaaaacgtaaatttgcctgt
Savignyi        ataaaataaggtttttgctttatt--------------------------attacattac
                 * ** *   ** .** ..*.* *                          ***   .*..

Robusta         tgttaagcggaggaaaaagtaatctgcgtgaatgcgaaaaacacgattttggaatcagcg
Savignyi        tgttaaaaataggagataggaaattgtg----tgtgatattcaggcagatcgtattggaa
                ******. . ****.* ** ** .**.*    **.** *  ** *    * * **..* .

Robusta         ccggcaatggtgtttgtaaa-taggggtggcatacgcgtttcgtagcgaaagagagaatg
Savignyi        actacgaagtaatttgtaaattgagggcagcatacgctgtccgggaagctaacgaga--g
                 * .*.* *  .******** *..***..********  *.** .. *  *. ****  *

Robusta         aggcgaaagtcgacagatgc--acgctccgatttatgagacaggaaccagtcgcgagggc
Savignyi        aagcaaaagtcgacagatgcgaaagctccgatttacgagataccaggcaat--------c
                *.**.***************  * ***********.****.*  *. **.*        *

Robusta         acggaggaaaaagaaccttactccaagacatgcgccgccttttttctttctgtcctagtc
Savignyi        tgggggagaaacgcaacttgttct-------------cgtcttttttcctcaatccgaac
                  **.*..*** * * ***..**.             * *.****.*..... .*... *

Robusta         aggaatactagagtatagaaggc-cacgcgtcgtggagtttaaaaccagcaag-ccagtg
Savignyi        gggaatactacagtatagaaggcacgcgcctcgctgagtttaaaaccagcgagcccagtg
                .********* ************ *.*** ***. ***************.** ******

Robusta         tctcaacggacacatcaatacagagacttctctcagtggacaactcggacgattcgccac
Savignyi        tctcaacggacacatcaatacccagcgttctctcagtggacagtcaggacg---------
                *********************  **  ***************... *****

Robusta         taacttggtggattcgtcccggacgacccatgg-gccgggtcccagcgcgctagttggcc
Savignyi        -aactttcagaactcgtctcgctcgacccaaggcgacgggtcccagcgcgctagtttact
                 *****   *.*.*****.**  ******* ** * ******************** .*.

Robusta         accatacagtgtagaatcagctagatcgtctctgcggatttcgcaaatagattgagttgg
Savignyi        gcctaatagcgactgcaacagtgacttgtctctgtgggcttcggaaatagattgactttg
                .**  *.**.*   .    . *.. *.*******.**..**** *********** ** *

Robusta         agatagcttcccgaccgggatttcgactaatttgcaatgttagttattaatcaaggtgac
Savignyi        agttttttt--------gactttcgtttaattttcattacgtagttataattaagacgat
                ** *  .**        *. ***** .****** ** *..  . *  ****.***..**.

Robusta         agtcaggagttaaagttaatttaccttttgaaagggcaaaaagattttcgaagtaaattg
Savignyi        cg--aagcattgcgattaat-----ttctatgaatgacaaacgttgtctggaataatctg
                 *  *.* .**. ..*****     **.*. .*. *  *** * * *..*.*.*** .**

Robusta         attccgttaattgtaactcttaaacgcaaaccgataaacgacgccattttgcttttcatt
Savignyi        tttc----gtaaataacaggaaaatatatattttttaatatcctagtcatatacgttatt
                 ***    .   .****    ***...* *..  * **.. * . .*. *.. . *.***

Robusta         gagaaactaaccatttaggcattctataattaaaattaaatgtttttaaaatctgtaact
Savignyi        acaag----------tggtagttttctccgttaaattgtttatttaaaaagacgttgctt
                . .*.          *.*  .**.* *   * *****.  *.***  ***. *  *. .*

Robusta         atctgtaataaactagaattatttatttcagtttaaatttttatttaaaacaataatttt
Savignyi        taatacagtaggctactaactaaatacttattcaaaagctatttttacattaattatttt
                   *..*.**..***  * .      .*.* *. *** .* * **** * .*** *****

Robusta         attatttcctttat--tcatcctaatgatattggtatagagaaaacgatgtttttatttc
Savignyi        ggtatgttagttaccattatatcgaaaacatttcaacaga-----------ttttgtttg
                . *** *.  ***.  *.** ...* .*.***   *.***           ****.***

Robusta         cataaaattttatttcagaaaagtcattttgttcaatttaaaaaattcctttattctaaa
Savignyi        aattaattatttgtgaagtaaaat------------------------------------
                 ** ** * **  *  ** ***.*

Robusta         aaaatgcccaaaacaagttctttatttcttaaaaattatcctaaataaaaaaatccacgt
Savignyi        ------------------------------------------------------------


Robusta         tttaataaaatgtataaaattgaaaactataaaaagatgacttatttttttaccctaacg
Savignyi        ---------atggatggacct----------------caacttccaggctcagcggtacg
                         *** **..* .*                ..**** .   .*.* *   ***

Robusta         tgatttttcaccagataccttaagtgttattttatttgtaagtaatatccaaatggcaac
Savignyi        tggttggatgcaaactagtctgattg---------------------aaccaatggcaa-
                **.**   ..* *. ** ..*.* **                       * ********

Robusta         aatcgcg
Savignyi        ------g
                      *
```

**Sequences of *cis-*regulatory regions used in reporter plasmid assays**

>oculata.Ebf -3654/+24

GGTTTGTGGTTTGATGGCTGAAATTCAAATCCAAAAGACAAAAAAGTAATTCTACTGATTTTTATCAGATTTAAAAAAATGACAATGAAGGAAAGAGTTTATTAATGCCACTATGTAGTTTGAAACATTTCGCTACATTAGGTTAAACATTCTACATTTGGCCAATAGCTAATTAACATATTTACTAACACTTTGTTGTTATTTTATAATATTGGCTATATAGTGGTCATTAAATTTGGAATCACTGGTCATTTAAATATGGATCTCGATTCAGAACGTTTTCACATTATAGTTTGCAGCTGTTGGTTTGCTATAAGTGCAAAACGAACATATAAGTTAAGCGCAGCAGGATAAACTTAAATATATTCTTGACGCTATATGTCGCAACAGAGCAATGAATGTTTATTACAGTGTTGGGTTATTGTATTATTTTGATTCAGTAATTAAATTTACCTTTCATTTAATTATGTCAAAGCAAAATGTAATTAAATACTTCCATCCATTTATCAAATAAAACCTTGAGTTGAAAACACGACCTCAGTGATTGGTTTAACGTAATTGTCAAAATAACATGCCAACAACAGTATGGTTTTAAAATTAACGTTTACTTATTAAAAAATTATACTATTTGTTTAATTTCTTAATACTCTCTTTGATTAAACTAAAAAACGGTTTTTGACACTTTTGTTTGCTTTTCATTTGTCTACCTGAAGCTCGGCCGATTACAATTGGCTGCCTGTGTTTAAACTCTGCTAATATTGTTGCTTTCATCATCATGGGGCATGTGGATTTGGTTTACACGTTGTAATAATAATATTTGTCTGATTTTTATTCATCTGATGTGGAGTTTTGGTTCCCATGACTGATTTTACGATGTGGTTGTAAGTCATTATTCCTGGGTCAGATTAAAAACCGACCGTACATTCGCAGAGGCATTTTAAGCATCGTTTTGTTCAAATCTGAATTTAGCCATCGTGTCCTTCGAATTTCATTTTGCCATAATTCGGCATTTAAAATGATCAGAAATGTCGCTAGAAACTCAAAACAATTTCAGTTCTCGAAATGATCTCGCGTGTTGATTTGCGTAAAAATTCCGACCAAGCTCACATTGTCTTTCAGAAATAAAAAAAAATATATTTCAACAAAAGCAAGATTGGCTGAAACTTCAATGTAAACCTGCAAAGCCACAAGTCGTGGAGACGCGTGGCGTTGCTGAGGTGTAGGCTAATTTCGAAAAGGTCATAATCGCATTTCGAAAATTTAATAAATGATAACGCGAAGCATCTTTGCGAATATATCGCTACGGAAATCTTTTTACGAATTCGCCGCAATTCCAATCCGTGACAAGTGACATCCGTAGATTAATTTTATTACGCCCACTGCACGCTTAACACAACATATGGACACCCGTCGCTAAACATACGCTCGGCTAAATCCGTGCAACGTTTTAATAAAAACTCGCTTTTCGTAGCATCCGTATAAATTTTCTGAAAAAAATTCACGAAATGAAATTTTATTTGTTTGTAATTGTAACAATGGTATTTCGCGTGGTAACAATTGCAATTGTGATAGTGTGCTTGCATGACTGCGTTTAATTCATTTATCTCCTTTAGTTACTGGGCTGGATATCTGCTGATGGGAGATTTTTAGCTCGATAGTGCGGCATATGGAGACGATGCAATTAATAATGCATGGCATATCGACTTTTCGGTTTTACTACGACGCGAGAGCGGAATGCATATCTACAAGGTATAAACTGACATGAATTCAATTCCGCCTCTGCTGCGAATCGCTGGTCAAACTTATCATCCTGTCCACCTGTTCGCGACATTTGCGATGTCCGCATCGTGTGACAGTTTACGTCATTTTCTTGACCAGTGACAATAAAATGACCTCAATGTATTTGTCATAATGGTCATTGTTTTTCTTTAAATCCAAATTTAATATCCAGCTATGATTTTATTTGCGATTATTTTTTGTTATTTCAGTAAATTAATCGAAAATTTTGTCATAGAAATCTTGGATTGAAATAAAATAGGTCCATATTTTGAGCATACCTTTTTTAATACTTCAATGAATAGAATACATTTTATTCATTAGAAACACAAATCTGGCGTTGACAATTGCAAAAAATTTAGTTCGATTTTCATCAACTCCAAAAGTTGGTTTTAACAAGAAGATTATTTTATGTTGCGAGTCGCGTAAACATGTGATGGTGAAACAAAATGGCGCTGCCTCCCCTAGCGTATAATTAATATAAAGAAATGATAATTAAGGTTCACTGAATCCCCGCGACATAATTATTTCAATATTTTCCACCTAGCGGGTAATTAAACAAAAAAGTTACACTTGGCCTGATAGAATAATCAGGGTTTGATTTTTTTTACATTTTTTTAAATACACTAATTAAATATTAGATGAAATAAATAAATAAATGCTATTGAGATTGTGTTTCAATACACATATTTGATTCCCTCAGGTTCTAATAACCGTAGATCAAGAAAAGCACGACAGATTAACAGAAAGTGAACTACAGGGAATAAGCTGTTGGTCATGTATATTCAATAAGATTTAATTCTAAATTACATTTATACAACAAAATTGTCCCAAACCTTTATATTAATCGACTAGCCAGAAATTGAGAATAACAAAGCCCATGAATAAACACAGTTTGTTTTATATCTTTACTGATCATAACATAAAAATATAATTTTACCAAAATGATATTTTGGCCGACAAATTTTGGCAAACAAAAGGGTTAACATTAATAATTGTATAAGAGTTTGATTAAATATTAAAACAATGTTCATTCAACTTTGGAAAATAAATCAATATTCCACGAAAAACATCTCAAATTAAAAGAAATGTTGATTTTACGCGTTTTCATTATGACCTCGTTACATTAAGGTTTGAAATTGGCATGAAATTGAAACGAGTTGTTGTGTAACAATCAGTGGAACAATCACAGCATTGTTTTGTTATTGTTCACTAATAATCATGTTGTCTTTGCACTGCGCTCTGTGACGACATATCGCAAGCGTAAAAAGCGACATATTGCATTAATTACTTGGTAATTCGTAACAGGGTCGTAAATAGATAACTGAAACATCCCGCTAGCAAAACACAATGACAGCACGCGACCGATTTTCGATGTTTATATTTCCACATAGTGACATATGTACGAGTGAGATGGCCAGCTCGATCATGAGAAAAAATATTTGTAGTTTGAAATAATGTTTAATTCACATCCAATAAAAAGAAATGGCGAAAAGGAAGAAGTTAGAGTTGCTGTGATGTGATTGGTCGATACCAGTCGAAGTAGATGAGTTATATAGGTTTTGTATCTCTATGGGTAGTCGATTCTAAGCAATGAAACCCAAGGGCATTGTACGCAGTCAATATGTTCGAAAGTCTTTGGAAGTTGATGTTCAGCATTCTCCTGAGGAATTAATTCAAGGTTCAGTTCAGCCCCTTAAGGGTTTCTCAGGTGATAATTTCCCAATGAGAAATAACTCGACCCAAACATGAAAATATTAAATTTACGTCGTGAAAAATCATTTAGCTTTTGCTTAAAAATGGAGTGGAAATTAAACAATTTTTTCACAGAGATTTTAAACAGTATTTTCATACAATTTTATATATATAGTGCATAAATGTCAAATTTACCAAGCCAACAGTAA

>occulta.Ebf -3659/+24

CATCACCAGTGTCGGAGAAGTGGATTCATCGACCTGCCATTTATATTTAGAGTTATAGGCCTATATGGCGATTAATAGACTTGGTTTTACCTTTTTATAATATAATATATTTCAATATTATTTTTAATTCCTAACTTTTATCCTAACTTGCTTCTGTTTATCTGTTAAGTGTAATATCGTTCTGCCTATAATACCCTGAATTACGGATCATAAATCGATTGATTGTTATTGAAACAATTAATTAAGTGCTTTATTTTGTTCCACTTAAAACCTCATCAAGCTTTGCATTTGAATTTTAAAAGTTTTCATGGTTTGGTGCTGGAATTCATACGCCAACGTAAAACATACCTCGTCACGCTATAGCAAAAAGTGCCTATCTCAGTTTTGGGGCATTTTGAGTAAATTGATACCAAAATCTTCAGAAAAACAAAGCGCATCGACTGGTGCAATTTCCTTTATCACATAGTAATCTTGTAAAACTATTGATGAAGAATTGTATTTTTTTTGTGAGATTTTTTAAGACAATTTTTTGCGGCATTTGAAATTTTGCATTATTTTTGTTTTAGCTTGTGGTTGATTTGAACAAAAAAGTGCCCATCTCAAACGTTGTTCAAGATATAGTGCCTATACATCTCATGCTATCTCAGTAAATAATAAATATTTCTTAATCAAAATTTATAGAGTCGTTTCTTTAAAATAGTGTAATTCATTAATGAAAAAAACATTTACGTATACCAATCTATTTTTTCGATTACCTCAGAACGACTAAAAATGAGATATAGGCAGTTTTCGTTTTAGCTTGGCGTTATGGTCAGTGGTTATGAACATAGCAAATAAATGCATTGGTCATTCAATTGGTCAGTTTATTCAAAACAAAATCTTTACATTGGTTTGCAGACGTAACACTAAAAATTAACGTTCTTAACACTGGGATAACTTTAATTTATTTCCAACGCTATATGTCGCACCAGAGCAATGGAAGGTTGTTTATTCTATTGTTTAGAGGCTATAATTAAATTTAACTTTTATTTGATTATGTCAAAGATAAATGTAATTAGATGCTTCCACCCATTCATCAAACAAAGAATTGAAAACACAACCTCAATGAATGGGGTTAAAATAATTGCCACAACAAATGTGTTAAAATATAAGATGGTCATGGCATTACATGATGGATTCCACGATGTGGTTGTAAGTCATTAAATGGACATAATAAAAACCGACAGTATTCGCAGACATCTTTATGCCACTGTTTCGTGAAAATTATTTTCATTGTACTTGGCTTTGATTTACTTTAAAATGCAAACCTGTAAAAAGATGTGATGTTGTCGAAGTGCAATTTCTAAAAGTCATAATCGCATTTGCTAAATTTTATAAATGCTTACACAAAGCATCTTTGCGAATATATCGCTACGGAAATCTTTTCGCCGCAATTTCAATCCGTGACAAGTGACATCCGTATAGATTAATTTTTTTTTACGCCCAATGCACGCTTAACACAACATATGGACACCCGTCGCTAAACATACGCTCGGCTAAATCCGTGCAACGTTATAATAAAATCTCGCTTTTCGTAGCAGACGTCGAAAATTTTAAAAACAATTGGAAATGAAATTTTATTTGATTGTATCTTTACAATGGTATTATGCTTGGTAACAATTGCAATTGTGATAGTGTATACTTGCATGGTTGCGTTTAATTCATTTATCTCCTTTAGATTTGCTGGGTTGGATATATCAGGGAGGTTTTTAGTTCGATAGTGCGGCATATGGAGAAGATATAATTAATAATGCATGGCGATATCGACTTTTTGGTTTTTACTACGACGGGAGAGCGAAAGCGGAATGCATACAAGGTATAAACTGACATGAATTCAATTCGCCTTTGCTGCGAATCGTTGGTCAAACTTAGCATCCTGTCCACCTGTTCGCGACATTTGTGATGTCCGTATCGCATGACAGTTTACGTCATTATCTTGACCAGTGACAATAAACTGTCATGAATGTATTTATCATAATGACCATTGTTTACCTTCTAATACAATACATCCAACAATTTTGTGATTATTTTTATATTTTAATAAATTAATCGAAAATTTTGTCACAGAAACATTGGATTAAAGTGAAATTGAAAAATAGGTTCTTAATTTTTGTTTAATTCTTTTGATAATGAATAATCCATTTTCTTCATCCGAAACACAAATGTGATCATTGCACGAAAAGCAGTTTGATCATCTCCAAAAGTTTGTTTGCTACTGGGAAAGATTGTTGTTGTTCTTGCGCAAACATGAGTTCGAACAAAATGGCGCTGACCCTTCAAGCGTGTAATTAAAATAAAGAATTGATGATTAAGGTTCATTAAATCCTCGCGACATCTCAAGCAATTATTTCAAAATTTCTCGACCTCACTTTTGTGGGTAACTAAAAAATATTTTGAAAAAATCGCGTTTTTTTTAAAGTACAATAATTAAATAAAATAAATAAATGCTATTGTTATTATGTTTCAATACACCTTAAGTTCTAATAATCATATATCAAGAAAAGCTCGACAGAATGTGAAGTATAGACGGATTTGAGCTGTTATGTTATTGTATATTCAATAAGATTTAATTCTAAATTACGTTTATACAACAAAATTGTCCTAAACCTTGGTTAATCCAATGGACAAGAAATTGAGAAATATCAAGTGTGATGTTAACAAGTATACAAGTAGATACACTACAGTATATTTTATTACTTTACTGGTGATAACATAAAATAAATTGTGCAAATAAAGGGTTAACATTAACATCGAAGTTTGATTAAATATTAAAACAATGTTCATTCAACTTTAGAAAATAAATCAATATTTAACGAAAAACATTCCAAATTAAAAGAAATGTTGATTTCATGATGACCTCATTGCATTAAGGTTGAAATTTGCATGAAATTGAAACGAGTTGTTATGTAACAATCGGTGGAACAATCACAACATTGTTTTGTTATTGTTCACTAATAATCGTATTGACTTAGCAGTGCGCTCTGTGACGACATAACGCAAGCGTAAAAAGTGACATATTGCATTAATTACTTGGTAATTCGTAACAGGGTCGTAAATAGATAACTGAAACATCCCGCTAGCAAAACACAATGACAGCACGCGACCGATTTTCGATGTTTATATTTCCACATAGTGACATATGTATCAGCAAAACGATGCCAGCTCGTTCATGAGAAAAAATATTTTCAATTTGAAATGTTTAATTCGCTTCCAATAAAAAGAAATGGCGAAAAGAAAGAAGTTATAGTTGCTGTGATGTGATTGGTCGATAGCAGTCGAAGGAGATGAGTTATATAGGTTTTATATCTCAGTGCCGTTGATTTTAAGCAATGAAACCCAAGGGCATTGTACGCAGTCAATATGTTCGAAAGCCTTGGAAGTGGATGCATTTCAATCTCTTGAGGAATTAATTCAAGGTTCAGTTCAGCCCCTTAAGGGTTTCTCAGGTGATAATTGCCCAATGAGGAATAACTCCACCCAGGAATGATGTGAAATATTTAAATTCCCAGTAAAAAAAATAATTTGGCTTTCTCATTAAAAAAAAGTGTGAAAAATAAACATTTCTTCACAGAGATTTTCAACAATATTTTCATACAATTTTATATTATATCGTGCATAAATGTCAAATTTACCAAGCCAACAGTAA

>Mocu-->Mocc (**swapped sequence in red**)

GGTTTGTGGTTTGATGGCTGAAATTCAAATCCAAAAGACAAAAAAGTAATTCTACTGATTTTTATCAGATTTAAAAAAATGACAATGAAGGAAAGAGTTTATTAATGCCACTATGTAGTTTGAAACATTTCGCTACATTAGGTTAAACATTCTACATTTGGCCAATAGCTAATTAACATATTTACTAACACTTTGTTGTTATTTTATAATATTGGCTATATAGTGGTCATTAAATTTGGAATCACTGGTCATTTAAATATGGATCTCGATTCAGAACGTTTTCACATTATAGTTTGCAGCTGTTGGTTTGCTATAAGTGCAAAACGAACATATAAGTTAAGCGCAGCAGGATAAACTTAAATATATTCTTGACGCTATATGTCGCAACAGAGCAATGAATGTTTATTACAGTGTTGGGTTATTGTATTATTTTGATTCAGTAATTAAATTTACCTTTCATTTAATTATGTCAAAGCAAAATGTAATTAAATACTTCCATCCATTTATCAAATAAAACCTTGAGTTGAAAACACGACCTCAGTGATTGGTTTAACGTAATTGTCAAAATAACATGCCAACAACAGTATGGTTTTAAAATTAACGTTTACTTATTAAAAAATTATACTATTTGTTTAATTTCTTAATACTCTCTTTGATTAAACTAAAAAACGGTTTTTGACACTTTTGTTTGCTTTTCATTTGTCTACCTGAAGCTCGGCCGATTACAATTGGCTGCCTGTGTTTAAACTCTGCTAATATTGTTGCTTTCATCATCATGGGGCATGTGGATTTGGTTTACACGTTGTAATAATAATATTTGTCTGATTTTTATTCATCTGATGTGGAGTTTTGGTTCCCATGACTGATTTTACGATGTGGTTGTAAGTCATTATTCCTGGGTCAGATTAAAAACCGACCGTACATTCGCAGAGGCATTTTAAGCATCGTTTTGTTCAAATCTGAATTTAGCCATCGTGTCCTTCGAATTTCATTTTGCCATAATTCGGCATTTAAAATGATCAGAAATGTCGCTAGAAACTCAAAACAATTTCAGTTCTCGAAATGATCTCGCGTGTTGATTTGCGTAAAAATTCCGACCAAGCTCACATTGTCTTTCAGAAATAAAAAAAAATATATTTCAACAAAAGCAAGATTGGCTGAAACTTCAATGTAAACCTGCAAAGCCACAAGTCGTGGAGACGCGTGGCGTTGCTGAGGTGTAGGCTAATTTCGAAAAGGTCATAATCGCATTTCGAAAATTTAATAAATGATAACGCGAAGCATCTTTGCGAATATATCGCTACGGAAATCTTTTTACGAATTCGCCGCAATTCCAATCCGTGACAAGTGACATCCGTAGATTAATTTTATTACGCCCACTGCACGCTTAACACAACATATGGACACCCGTCGCTAAACATACGCTCGGCTAAATCCGTGCAACGTTTTAATAAAAACTCGCTTTTCGTAGCATCCGTATAAATTTTCTGAAAAAAATTCACGAAATGAAATTTTATTTGTTTGTAATTGTAACAATGGTATTTCGCGTGGTAACAATTGCAATTGTGATAGTGTGCTTGCATGACTGCGTTTAATTCATTTATCTCCTTTAGTTACTGGGCTGGATATCTGCTGATGGGAGATTTTTAGCTCGATAGTGCGGCATATGGAGACGATGCAATTAATAATGCATGGCATATCGACTTTTCGGTTTTACTACGACGCGAGAGCGGAATGCATATCTACAAGGTATAAACTGACATGAATTCAATTCCGCCTCTGCTGCGAATCGCTGGTCAAACTTATCATCCTGTCCACCTGTTCGCGACATTTGCGATGTCCGCATCGTGTGACAGTTTACGTCATTTTCTTGACCAGTGACAATAAAATGACCTCAATGTATTTGTCATAATGGTCATTGTTTTTCTTTAAATCCAAATTTAATATCCAGCTATGATTTTATTTGCGATTATTTTTTGTTATTTCAGTAAATTAATCGAAAATTTTGTCATAGAAATCTTGGATTGAAATAAAATAGGTCCATATTTTGAGCATACCTTTTTTAATACTTCAATGAATAGAATACATTTTATTCATTAGAAACACAAATCTGGCGTTGACAATTGCAAAAAATTTAGTTCGATTTTCATCAACTCCAAAAGTTGGTTTTAACAAGAAGATTATTTTATGTTGCGAGTCGCGTAAACATGTGATGGTGAAACAAAATGGCGCTGCCTCCCCTAGCGTATAATTAATATAAAGAAATGATAATTAAGGTTCACTGAATCCCCGCGACATAATTATTTCAATATTTTCCACCTAGCGGGTAATTAAACAAAAAAGTTACACTTGGCCTGATAGAATAATCAGGGTTTGATTTTTTTTACATTTTTTTAAATACACTAATTAAATATTAGATGAAATAAATAAATAAATGCTATTGAGATTGTGTTTCAATACACATATTTGATTCCCTCAGGTTCTAATAACCGTAGATCAAGAAAAGCACGACAGATTAACAGAAAGTGAACTACAGGGAATAAGCTGTTGGTCATGTATATTCAATAAGATTTAATTCTAAATTACATTTATACAACAAAATTGTCCCAAACCTTTATATTAATCGACTAGCCAGAAATTGAGAATAACAAAGCCCATGAATAAACACAGTTTGTTTTATATCTTTACTGATCATAACATAAAAATATAATTTTACCAAAATGATATTTTGGCCGACAAATTTTGGCAAACAAAAGGGTTAACATTAATAATTGTATAAGAGTTTGATTAAATATTAAAACAATGTTCATTCAACTTTGGAAAATAAATCAATATTCCACGAAAAACATCTCAAATTAAAAGAAATGTTGATTTTACGCGTTTTCATTATGACCTCGTTACATTAAGGTTTGAAATTGGCATGAAATTGAAACGAGTTGTTGTGTAACAATCAGTGGAACAATCACAGCATTGTTTTGTTATTGTTCACTAATAATCATGTTGTCTTTGCACTGCGCTCTGTGACGACATATCGCAAGCGTAAAAAGCGACATATTGCATTAATTACTTGGTAATTCGTAACAGGGTCGTAAATAGATAACTGAAACATCCCGCTAGCAAAACACAATGACAGCACGCGACCGATTTTCGATGTTTATATTTCCACATAGTGACATATGTA**tcagcaaaacgat**GCCAGCTCGATCATGAGAAAAAATATTTGTAGTTTGAAATAATGTTTAATTCACATCCAATAAAAAGAAATGGCGAAAAGGAAGAAGTTAGAGTTGCTGTGATGTGATTGGTCGATACCAGTCGAAGTAGATGAGTTATATAGGTTTTGTATCTCTATGGGTAGTCGATTCTAAGCAATGAAACCCAAGGGCATTGTACGCAGTCAATATGTTCGAAAGTCTTTGGAAGTTGATGTTCAGCATTCTCCTGAGGAATTAATTCAAGGTTCAGTTCAGCCCCTTAAGGGTTTCTCAGGTGATAATTTCCCAATGAGAAATAACTCGACCCAAACATGAAAATATTAAATTTACGTCGTGAAAAATCATTTAGCTTTTGCTTAAAAATGGAGTGGAAATTAAACAATTTTTTCACAGAGATTTTAAACAGTATTTTCATACAATTTTATATATATAGTGCATAAATGTCAAATTTACCAAGCCAACAGTAA

>Mocc-->Mocu (**swapped sequence in blue**)

CATCACCAGTGTCGGAGAAGTGGATTCATCGACCTGCCATTTATATTTAGAGTTATAGGCCTATATGGCGATTAATAGACTTGGTTTTACCTTTTTATAATATAATATATTTCAATATTATTTTTAATTCCTAACTTTTATCCTAACTTGCTTCTGTTTATCTGTTAAGTGTAATATCGTTCTGCCTATAATACCCTGAATTACGGATCATAAATCGATTGATTGTTATTGAAACAATTAATTAAGTGCTTTATTTTGTTCCACTTAAAACCTCATCAAGCTTTGCATTTGAATTTTAAAAGTTTTCATGGTTTGGTGCTGGAATTCATACGCCAACGTAAAACATACCTCGTCACGCTATAGCAAAAAGTGCCTATCTCAGTTTTGGGGCATTTTGAGTAAATTGATACCAAAATCTTCAGAAAAACAAAGCGCATCGACTGGTGCAATTTCCTTTATCACATAGTAATCTTGTAAAACTATTGATGAAGAATTGTATTTTTTTTGTGAGATTTTTTAAGACAATTTTTTGCGGCATTTGAAATTTTGCATTATTTTTGTTTTAGCTTGTGGTTGATTTGAACAAAAAAGTGCCCATCTCAAACGTTGTTCAAGATATAGTGCCTATACATCTCATGCTATCTCAGTAAATAATAAATATTTCTTAATCAAAATTTATAGAGTCGTTTCTTTAAAATAGTGTAATTCATTAATGAAAAAAACATTTACGTATACCAATCTATTTTTTCGATTACCTCAGAACGACTAAAAATGAGATATAGGCAGTTTTCGTTTTAGCTTGGCGTTATGGTCAGTGGTTATGAACATAGCAAATAAATGCATTGGTCATTCAATTGGTCAGTTTATTCAAAACAAAATCTTTACATTGGTTTGCAGACGTAACACTAAAAATTAACGTTCTTAACACTGGGATAACTTTAATTTATTTCCAACGCTATATGTCGCACCAGAGCAATGGAAGGTTGTTTATTCTATTGTTTAGAGGCTATAATTAAATTTAACTTTTATTTGATTATGTCAAAGATAAATGTAATTAGATGCTTCCACCCATTCATCAAACAAAGAATTGAAAACACAACCTCAATGAATGGGGTTAAAATAATTGCCACAACAAATGTGTTAAAATATAAGATGGTCATGGCATTACATGATGGATTCCACGATGTGGTTGTAAGTCATTAAATGGACATAATAAAAACCGACAGTATTCGCAGACATCTTTATGCCACTGTTTCGTGAAAATTATTTTCATTGTACTTGGCTTTGATTTACTTTAAAATGCAAACCTGTAAAAAGATGTGATGTTGTCGAAGTGCAATTTCTAAAAGTCATAATCGCATTTGCTAAATTTTATAAATGCTTACACAAAGCATCTTTGCGAATATATCGCTACGGAAATCTTTTCGCCGCAATTTCAATCCGTGACAAGTGACATCCGTATAGATTAATTTTTTTTTACGCCCAATGCACGCTTAACACAACATATGGACACCCGTCGCTAAACATACGCTCGGCTAAATCCGTGCAACGTTATAATAAAATCTCGCTTTTCGTAGCAGACGTCGAAAATTTTAAAAACAATTGGAAATGAAATTTTATTTGATTGTATCTTTACAATGGTATTATGCTTGGTAACAATTGCAATTGTGATAGTGTATACTTGCATGGTTGCGTTTAATTCATTTATCTCCTTTAGATTTGCTGGGTTGGATATATCAGGGAGGTTTTTAGTTCGATAGTGCGGCATATGGAGAAGATATAATTAATAATGCATGGCGATATCGACTTTTTGGTTTTTACTACGACGGGAGAGCGAAAGCGGAATGCATACAAGGTATAAACTGACATGAATTCAATTCGCCTTTGCTGCGAATCGTTGGTCAAACTTAGCATCCTGTCCACCTGTTCGCGACATTTGTGATGTCCGTATCGCATGACAGTTTACGTCATTATCTTGACCAGTGACAATAAACTGTCATGAATGTATTTATCATAATGACCATTGTTTACCTTCTAATACAATACATCCAACAATTTTGTGATTATTTTTATATTTTAATAAATTAATCGAAAATTTTGTCACAGAAACATTGGATTAAAGTGAAATTGAAAAATAGGTTCTTAATTTTTGTTTAATTCTTTTGATAATGAATAATCCATTTTCTTCATCCGAAACACAAATGTGATCATTGCACGAAAAGCAGTTTGATCATCTCCAAAAGTTTGTTTGCTACTGGGAAAGATTGTTGTTGTTCTTGCGCAAACATGAGTTCGAACAAAATGGCGCTGACCCTTCAAGCGTGTAATTAAAATAAAGAATTGATGATTAAGGTTCATTAAATCCTCGCGACATCTCAAGCAATTATTTCAAAATTTCTCGACCTCACTTTTGTGGGTAACTAAAAAATATTTTGAAAAAATCGCGTTTTTTTTAAAGTACAATAATTAAATAAAATAAATAAATGCTATTGTTATTATGTTTCAATACACCTTAAGTTCTAATAATCATATATCAAGAAAAGCTCGACAGAATGTGAAGTATAGACGGATTTGAGCTGTTATGTTATTGTATATTCAATAAGATTTAATTCTAAATTACGTTTATACAACAAAATTGTCCTAAACCTTGGTTAATCCAATGGACAAGAAATTGAGAAATATCAAGTGTGATGTTAACAAGTATACAAGTAGATACACTACAGTATATTTTATTACTTTACTGGTGATAACATAAAATAAATTGTGCAAATAAAGGGTTAACATTAACATCGAAGTTTGATTAAATATTAAAACAATGTTCATTCAACTTTAGAAAATAAATCAATATTTAACGAAAAACATTCCAAATTAAAAGAAATGTTGATTTCATGATGACCTCATTGCATTAAGGTTGAAATTTGCATGAAATTGAAACGAGTTGTTATGTAACAATCGGTGGAACAATCACAACATTGTTTTGTTATTGTTCACTAATAATCGTATTGACTTAGCAGTGCGCTCTGTGACGACATAACGCAAGCGTAAAAAGTGACATATTGCATTAATTACTTGGTAATTCGTAACAGGGTCGTAAATAGATAACTGAAACATCCCGCTAGCAAAACACAATGACAGCACGCGACCGATTTTCGATGTTTATATTTCCACATAGTGACATATGTA**cgagtgagatg**GCCAGCTCGTTCATGAGAAAAAATATTTTCAATTTGAAATGTTTAATTCGCTTCCAATAAAAAGAAATGGCGAAAAGAAAGAAGTTATAGTTGCTGTGATGTGATTGGTCGATAGCAGTCGAAGGAGATGAGTTATATAGGTTTTATATCTCAGTGCCGTTGATTTTAAGCAATGAAACCCAAGGGCATTGTACGCAGTCAATATGTTCGAAAGCCTTGGAAGTGGATGCATTTCAATCTCTTGAGGAATTAATTCAAGGTTCAGTTCAGCCCCTTAAGGGTTTCTCAGGTGATAATTGCCCAATGAGGAATAACTCCACCCAGGAATGATGTGAAATATTTAAATTCCCAGTAAAAAAAATAATTTGGCTTTCTCATTAAAAAAAAGTGTGAAAAATAAACATTTCTTCACAGAGATTTTCAACAATATTTTCATACAATTTTATATTATATCGTGCATAAATGTCAAATTTACCAAGCCAACAGTAA
